# DART: Deep learning for the Analysis and Reconstruction of Transcriptional dynamics from live-cell imaging data

**DOI:** 10.1101/2025.09.02.673499

**Authors:** Muhan Ma, Ramon Grima

## Abstract

Transcriptional bursting, characterized by stochastic switching between promoter states, underlies cell-to-cell variability in gene expression. Accurately inferring promoter activity from live-cell imaging data remains challenging because the fluorescence signal at any given point is influenced by the history of promoter states. Here, we present DART (Deep learning for the Analysis and Reconstruction of Transcriptional dynamics), a deep learning framework that infers promoter onand off-states from fluorescence intensity traces, enabling the estimation of activation and inactivation rates and the selection of the most appropriate promoter-switching model. DART utilizes a neural network architecture that combines convolutional neural networks and long short-term memory layers to binarize fluorescence traces. Using extensive synthetic datasets spanning a wide range of transcriptional bursting levels, we demonstrate that DART outperforms current binarization methods, including conventional and augmented hidden Markov models, in both accuracy and robustness. Furthermore, a reanalysis of published experimental data using DART reveals a strong linear coupling between activation and inactivation rates, contradicting previous claims of independence. Our approach provides a powerful and generalizable tool for quantitative analysis of transcriptional kinetics from live-cell imaging data.

## 1 Introduction

Biochemical dynamics is fundamentally stochastic due to the inherently probabilistic nature of molecular interactions [1]. Specifically, the timing of the reaction events and the order in which successive reactions occur are random, which in turn leads to stochasticity in the number of molecules, a phenomenon often called intrinsic noise [2]. This partially accounts for the cell-to-cell variation in the number of mRNA observed by single-molecule fluorescence in situ hybridization (smFISH). The rest of the variability is principally due to the variation of rate parameter values across cells (extrinsic noise); this can be attributed to differences between cells due to size [3, 4], cell-cycle phase [5–7], the extent of macromolecular crowding, and the concentrations and locations of polymerases and DNA-binding proteins in the nucleus [8, 9] at the time of measurement.

By fitting snapshot measurements of the mRNA distribution (or of its moments) obtained using smFISH and single-cell sequencing (scRNA-seq) to mechanistic stochastic models of gene expression, several studies have attempted to reconstruct the promoter switching characteristics of thousands of genes in eukaryotic cells [7, 10–18]. In principle, by these methods, one can deduce the number of promoter states, their connectivity, and parameter values for the rates of switching between different states. A major difficulty with this approach is that mRNA distributions are strongly affected by extrinsic and technical noise [15, 19–22] and therefore inference of the number of gene states and their connectivity from snapshot data is physically meaningful to the extent that these noise sources can be properly accounted for within the mechanistic modeling framework.

It is also the case that many measurements of cellular mRNA copy numbers are of mature mRNA which are strongly affected by processes downstream of transcription; unless the models also account for this noise, it can significantly affect the accuracy of transcriptional parameter inference [6, 18].

Live-cell imaging provides an ideal way to bypass these problems by visualizing in real-time nascent RNA production in individual cells [23–28]. The commonest used technique of this type is the MS2/MCP system [29]. This works by inserting RNA stem-loop sequences (MS2) into the gene of interest, which are specifically bound by fluorescently labeled RNA-binding coat proteins MS2 (MCP) when RNA is transcribed (Fig. 1A). In principle, the peaks of the fluorescence signal correspond to gene activity and the troughs to gene inactivity, thus opening up the possibility of directly observing the promoter switching dynamics. However, typical measured time series do not closely resemble a set of square pulses with stochastic durations, which is what is expected from the switching between active and inactive promoter states. Consequently, methods are needed to binarize the fluorescence signal.

**Fig. 1.**
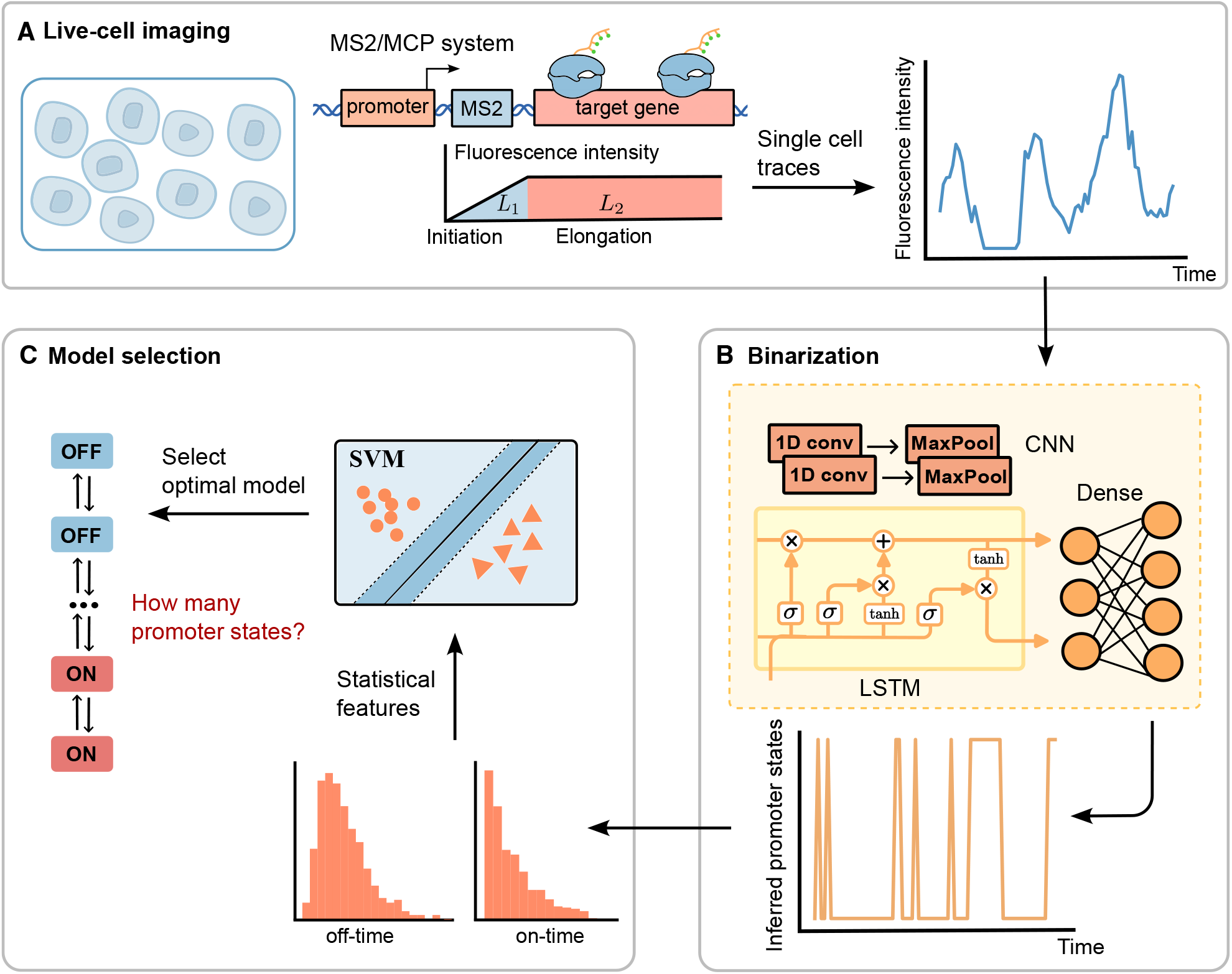
Schematic of DART. (A) The MS2/MCP system enables real-time visualization of nascent mRNA molecules, generating fluorescence signals at the single-cell level. Here *L*represents the length of the MS2 sequence, and *L*_2_ denotes the gene length. (B) A deep learning model comprising two convolutional neural network (CNN) layers, one long short-term memory (LSTM) layer, and one dense layer was employed to infer promoter states (on and off) from single-cell fluorescence signal traces. (C) Distributions of on- and off-times were extracted from the binarized promoter state traces, and statistical features derived from these distributions were used as inputs to a support vector machine (SVM) model to perform model selection and determine the optimal number of promoter on and off states.

One of the simplest methods to achieve this is a two-state Hidden Markov Model (HMM) [27, 30–33]. The output is a binary trace that represents the activity and inactivity of a gene. The assumption behind this approach is that the instantaneous fluorescence intensity depends on a hidden state (which can be active or inactive) whose duration is exponentially distributed. Alternatively, regions of transcriptional activity can be identified as those parts of the time trace where the signal or a suitably smoothened version of the signal exceeds a user-defined-threshold [32, 34]. Some have argued that these simple methods may give incorrect results because the fluorescence at any given time point is influenced, to some extent, by both the current and preceding promoter states. This memory effect can be understood as follows. When the promoter transitions to an active state, RNA Polymerase II (Pol II) begins elongation along the gene body, generating a fluorescent signal via binding of MCP-fluorescent proteins to nascent mRNA. Upon promoter inactivation, this fluorescence does not stop immediately, as Pol II molecules already engaged in transcription continue elongating, and the partially synthesized mRNA remains bound by MCP-fluorescent proteins [35, 36]. Augmented HMMs such as cpHMM [36] (which builds on an earlier approach by [35]), a much faster version of cpHMM called burstInfer [28] and other methods [37] have been designed to output a binarization of the fluorescence signal that is partly corrected for memory effects.

In this paper, we introduce DART (Deep learning for Analysis and Reconstruction of Transcriptional dynamics), a novel deep learning-based framework for the analysis of live-cell imaging data. DART combines convolutional and recurrent neural network architectures to robustly infer gene activity states from noisy fluorescence traces and leverages machine learning classifiers to distinguish among promoter models with varying numbers of states (Fig. 1). By training on a large number of synthetic datasets generated using stochastic simulations that capture a wide range of biologically realistic transcriptional dynamics, we show that DART achieves high accuracy in both state inference and model selection, outperforming existing methods under both idealized and experimentally realistic conditions. Furthermore, a re-analysis of published experimental data using DART reveals a linear coupling between activation and inactivation rates, contradicting previous claims of independence.

## 2 Results

### 2.1 DART outperforms standard binarization methods on optimally sampled synthetic data

We considered three commonly used binarization approaches: a two-state Hidden Markov Model (HMM), and thresholding methods based on moving average (MA) and Savitzky-Golay (SG) smoothing (for implementation details of these methods, see Materials and Methods, Section 4.1). To systematically assess the accuracy of commonly used binarization methods in capturing transcriptional bursting dynamics, we used these methods to infer the promoter state—on (transcriptionally active) or off (inactive)—from fluorescence intensity time series data generated by simulations of a gene expression model performed using the delay stochastic simulation algorithm (SSA). These simulations mimic the data generated in a live-cell imaging experiment, assuming that intrinsic noise from the gene of interest can be explained by a model with *L* off-states and *N − L* on-states (for details see Materials and Methods, Section 4.2.1). The existence of multiple on- or off-states for many eukaryotic genes is suggested by many previous studies [38–45]. This approach offers a more accurate model of gene expression dynamics compared to the classical two-state telegraph model [46], which assumes a single on-state and a single off-state. Since the true promoter state at each time point is known from the simulations, we directly compared the states inferred by the binarization methods to the ground truth to assess each method’s accuracy.

For this comparison, we use data from 1800 biologically realistic rate parameter sets of the gene expression model with three on- and three off-states (*L* = 3, *N* = 6), and assume idealized observation conditions, i.e. where the time between successive measurements and the total cell number are optimally selected for each parameter set of the gene expression model such that the fluorescence intensity time series have sufficient statistical information about the underlying promoter switching dynamics. Further technical details on the parameter ranges and other pertinent details can be found in Materials and Methods, Section 4.2.2 and 4.5.1.

The quantitative evaluation of the accuracy of the baseline binarization methods was based on the following three metrics calculated from the binarized time series: (1) the accuracy of the binary vector *α*, defined as the proportion of correctly classified promoter states in time; (2) the estimate of the mean on-time; and (3) the estimate of the mean off-time. We define Λ = τ/⟨*t*_off_⟩ where *τ* is the sum of the elongation and termination times and ⟨*t*_off_⟩ is the true mean time spent in the off states (Materials and Methods, Section 4.2.2). The results for the accuracy metrics, grouped by Λ, are shown in the first three rows of Fig. 2. We find that the three binarization methods have comparable accuracy: ⟨*α*⟩ ≈ 0.69 when Λ > 0.5 and ⟨*α*⟩ ≈ 0.90 when Λ *≤* 0.5. Note that the angled brackets denote an average over parameter sets. Thus the performance of the three baseline methods improves as Λ decreases. The explanation for this is as follows. After the promoter switches off, some RNAP molecules remain attached to the gene and take approximately time *τ* to elongate and detach, leading the baseline methods to erroneously assign this period as transcriptionally active (for an illustration see Fig. 3A); this error is particularly large when the time spent in the off states is comparable to or smaller than *τ*, i.e. large Λ (compare the first two rows of Fig. 3B and C). A different measure of the accuracy of the baseline methods is obtained from the plots of the true and predicted mean on-times, and of the true and predicted mean off-times (true on- and off-times computed directly from the synthetic traces generated by the simulations). The points in the mean-on times plots shown in the second column of Fig. 2 (each point corresponds to a different sampled parameter set) typically fall above the line *y* = *x* due to the overestimation of the duration of the on-state. Similar is observed for the mean off-times plots shown in the third column of Fig. 2; the overestimation of the mean off-time stems from the difficulty in detecting short, transient off-periods that occur between closely spaced on-periods, thereby inflating estimates of the mean off-times. The sizable overestimation in these plots is also evident from the small coefficient of determination (*R*^2^): 0.16 *−* 0.20 for the plots of the mean on-times and 0.35 *−* 0.50 for the plots of the mean off-times.

**Fig. 2.**
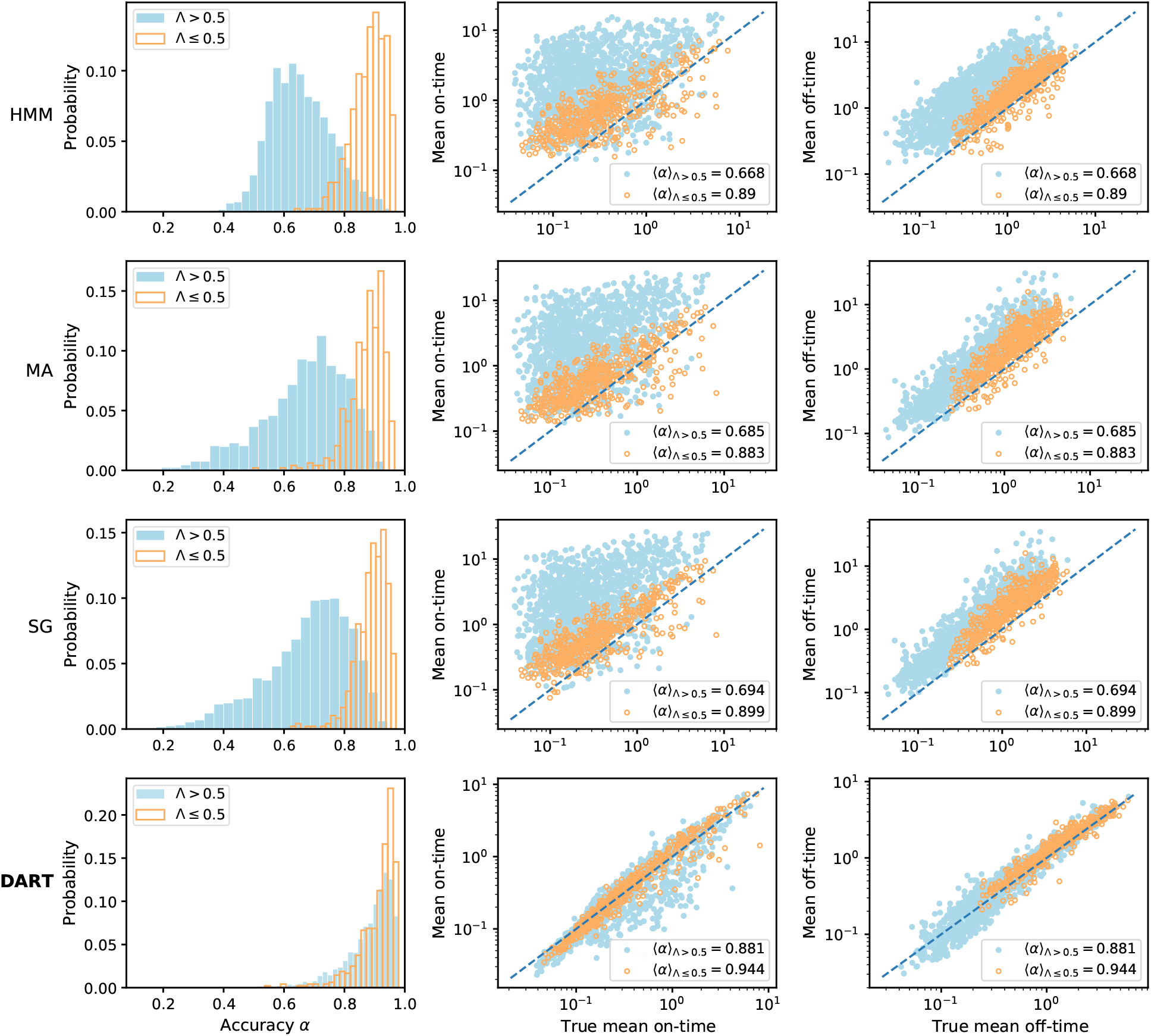
Binarization accuracy of common binarization methods (HMM, MA and SG) and of the deep learning method, DART. The first column shows the distribution of the proportion of correctly classified promoter states in time, i.e. the accuracy *α*. The second and third columns shows plots of the predicted and true mean-on times, and the predicted and true mean off-times, respectively (the time unit is minutes). In all three cases, these statistics are grouped by Λ which is the ratio of sum of the elongation and termination time to the true mean off-time. The ground truth traces of promoter-state values and the corresponding true means of the on- and off-time distributions are obtained from stochastic simulations using the SSA. The average accuracy ⟨*α*⟩ is computed over all parameter sets. Note that the DART results shown correspond to the random seed that produced median performance for the given metric, based on the posterior distribution from 100 independent runs (Supplementary Text S2 and Supplementary Fig. S2).

**Fig. 3.**
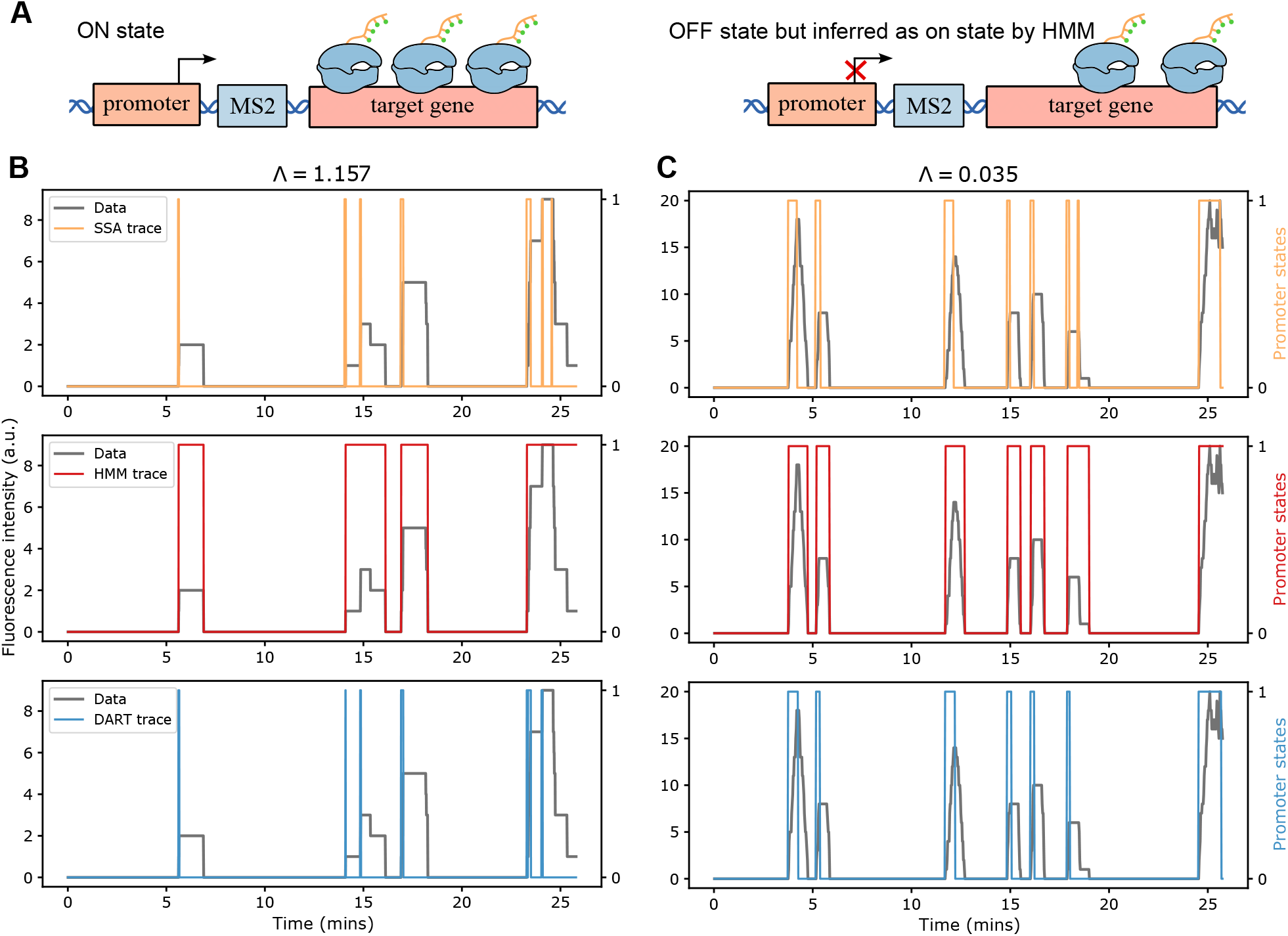
Comparison of the binarized traces output by HMM and DART. (A) Schematic illustrating why common baseline methods such as the HMM can mischaracterize states. This occurs when a promoter that is in the on state (left) switches off (right) but there is still residual fluorescence due to bound RNAPs that are yet to detach from the gene. (B)(C) Comparison of the time series of ground-truth promoter states from the SSA (orange trace) with that reconstructed by HMM (red trace) and DART (blue trace) for high and low values of Λ. The fluorescence signal is shown in grey. The binarized traces reconstructed by DART closely match the ground-truth states across all values of Λ, whereas those inferred by HMM are only reasonably accurate at low Λ.

Since the accuracy of these methods is far from satisfactory, we devised DART, a deep learning model that also binarizes fluorescence time series data. DART is initially trained on a large set of stochastic simulations to learn a mapping from fluorescence intensity time series to the corresponding ground-truth promoter state sequences. Once trained, it can infer binarized promoter state traces from new, previously unseen time series of the same type. For an illustration of how DART works see Fig. 1B. For technical details of DART and its training see Materials and Methods, Section 4.3 and 4.5.2. In the last row of Fig. 2 we compare the performance of DART with the three baseline methods (for the same input synthetic fluorescence intensity data). We find that DART consistently outperforms the three conventional binarization methods across all three evaluation metrics. In particular, the mean binary vector accuracy ⟨*α*⟩ is *≈* 0.88 for Λ *>* 0.5 and *≈* 0.94 for Λ *≤* 0.5. In addition, the true and predicted mean on-times in many cases fall on or close to the line *y* = *x* (same for the mean off-times) — in fact the values of *R*^2^ are 0.82 and 0.94 for the mean on-time and mean off-time plots, respectively. These results indicate strong agreement of the predicted promoter states by DART and the ground-truth promoter states across diverse transcriptional bursting regimes. The improvement offered by DART is most visually apparent when comparing the ground-truth state time series with the trajectories inferred by DART and HMM at both small and large Λ values (Fig. 3B,C). Notably, we found that DART also accurately infers binarized promoter states from synthetic data generated by types of stochastic gene expression models that it has not seen in its training (see Supplementary Text S1 and Supplementary Fig. S1 for details).

### 2.2 DART is robust under realistic experimental sampling protocols

To evaluate DART under more experimentally realistic sampling protocols, we generated noisy synthetic data based on experimental settings reported in previous studies [26, 47–50]. We simulated fluorescence intensity data for 300 cells, with each single-cell trace spanning 30 minutes and sampled every 6 seconds and we added log-normal noise to the synthetic fluorescence data to mimic background noise [6]. Further technical details can be found in Materials and Methods Section 4.5.3.

In Fig. 4 we show an evaluation of DART’s accuracy for low and high transcriptional bursting conditions, with technical noise having a coefficient of variation (CV) of 0% and 5%. For high transcriptional bursting, DART’s performance is practically independent of the size of technical noise with an accuracy of ⟨*α*⟩ ≈ 0.95 (first row of Fig. 4). In addition, we found that DART successfully reconstructed the full distributions of on- and off-times from synthetic fluorescence intensity data — the P-P plots show that cumulative distribution functions (CDFs) of the true on- and off-times, generated by the SSA, closely match those estimated by DART, with points aligning along the diagonal (second row of Fig. 4). For low transcriptional bursting (third and fourth rows of Fig. 4), we find similar results. Overall, DART’s accuracy appears to be primarily influenced by the level of transcriptional bursting (⟨*α*⟩ ≈ 0.95 for high bursting vs. ⟨*α*⟩ ≈ 0.85 for low bursting), rather than by the magnitude of technical noise. For an evaluation of DART’s performance for an intermediate level of transcriptional bursting and with higher levels of technical noise, see Supplementary Fig. S3 and Fig. S4, respectively.

**Fig. 4.**
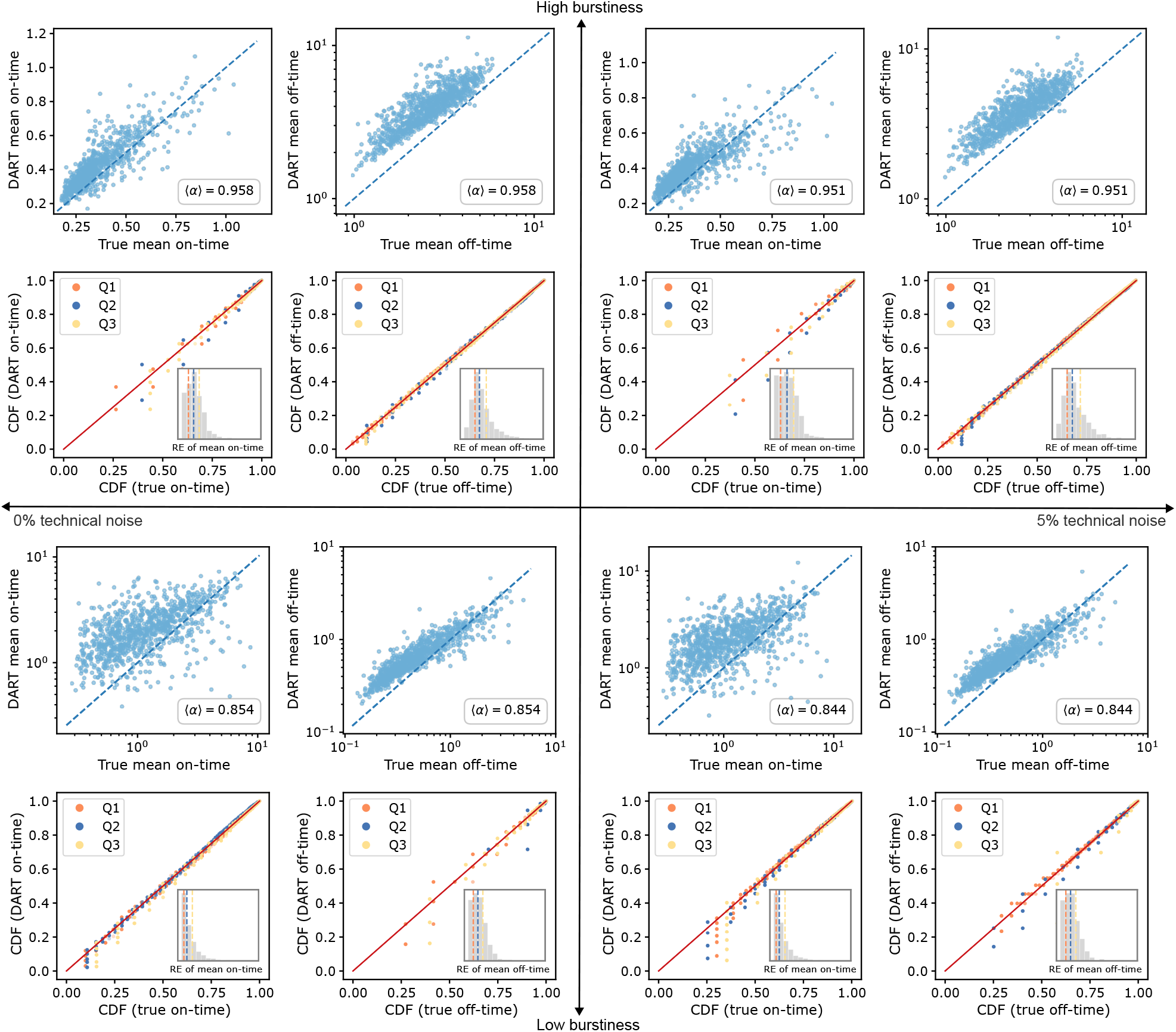
Performance of DART using synthetic data collected under realistic experimental sampling conditions. The four panels represent different combinations of the size of transcriptional burstiness and the size of technical (log-normal distributed) noise: top left—high burstiness with no technical noise; top right—high burstiness with 5% technical noise; bottom left—low burstiness with no technical noise; bottom right—low burstiness with 5% technical noise. In each case, the first row compares the true mean on- and off-times obtained from the SSA with the estimates from DART. The second row shows P–P plots comparing the true CDFs, obtained via the SSA, with those generated by DART, for three parameter sets corresponding to the 0.25 (Q1), 0.5 (Q2), and 0.75 (Q3) quantiles of the relative error (RE) distributions of the mean on- and off-time estimates (RE distributions calculated from the RE of all sampled parameters are shown in the insets). Note that the coarse appearance of the P-P plots of on-time (off-time) for high (low) burstiness results from limited data on the on-periods (off-periods) in these cases.

Next, we compared the performance of DART with a recently developed variant of the HMM method, burstInfer [28], which is a computationally optimized version of the cpHMM approach proposed in [36]. For reference, the basic HMM is also included here as a baseline. Other baseline methods assessed in the previous section are omitted here, as their performance was shown to be comparable to HMM (Fig. 2). As mentioned in the Introduction, cpHMM (and thus burstInfer) is a corrected version of HMM because it accounts, to some extent, for the memory in the fluorescent intensity signal due to bound RNAPs when the promoter is off. Due to the high computational cost of burstInfer, with inference for long genes that require approximately 14 hours and for short genes around 40 minutes for a single parameter set — estimated on an Intel® Xeon® Silver 4314 (2.40 GHz) compute node with 128 GB RAM — we restricted our comparison to a small number of parameter sets. Specifically, we selected 10 parameter sets near the first quartile (Q1) and 10 near the third quartile (Q3) of DART’s distribution of the promoter state accuracy *α*. The comparison was carried out under two levels of technical noise, with a CV of 5% and 20%. As shown in Fig. 5A, DART consistently outperformed both the burstInfer approach and the basic HMM approach for all levels of transcriptional burstiness and of technical noise. The limitations of burstInfer are particularly evident at low transcriptional burstiness levels, where it fails to capture a substantial fraction of promoter activation events, as shown in Fig. 5B. We also compared results using a synthetic fluorescent intensity trace with technical noise characterized by a CV of 30%; however, burstInfer failed for the majority of parameter sets, frequently inferring only a single promoter state, while DART achieved mean accuracies of 0.92, 0.82, 0.81 for high, medium, and low levels of transcriptional bursting, respectively.

**Fig. 5.**
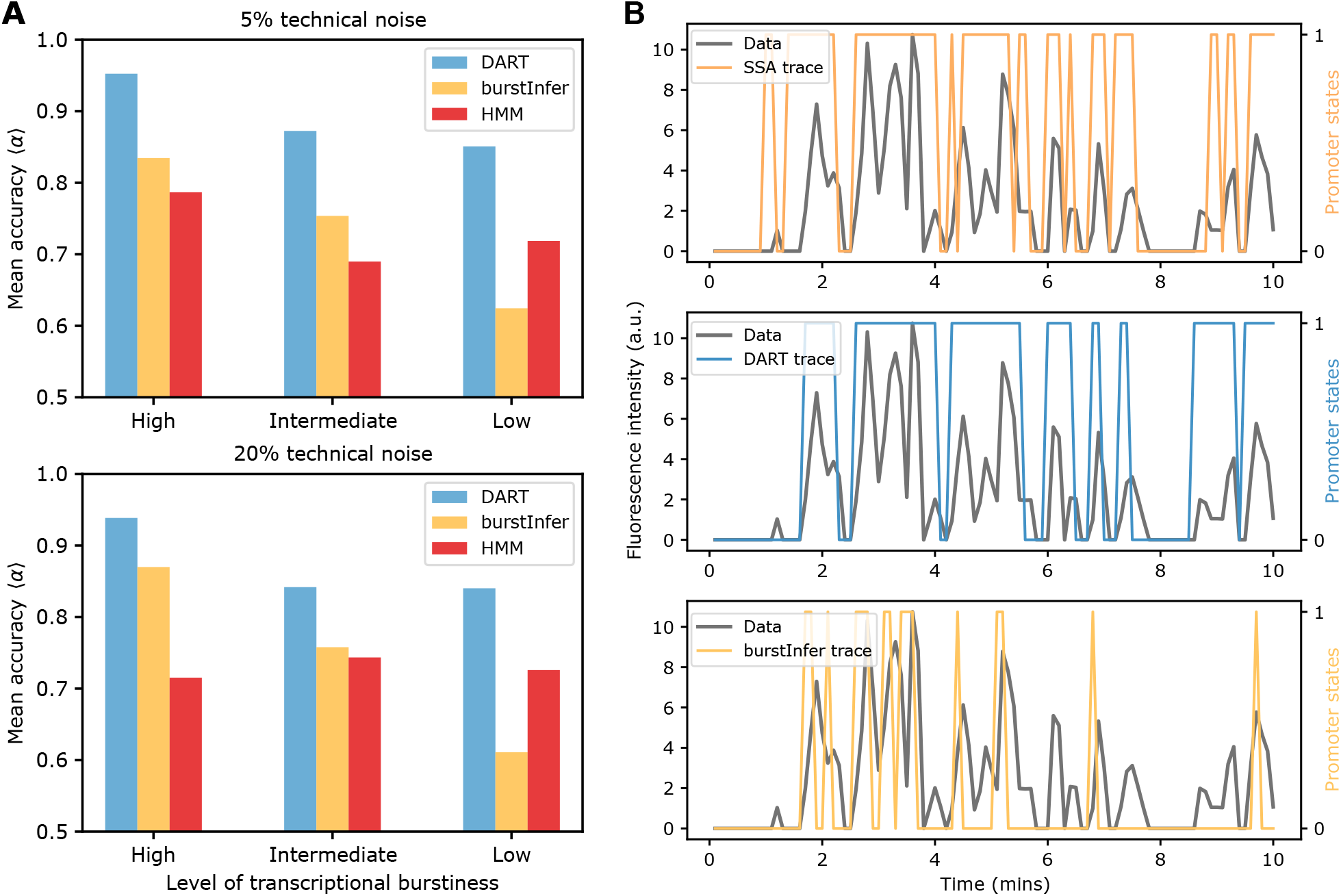
Comparison of DART, HMM and burstInfer (an augmented type of HMM to correct for memory effects in the intensity trace) using synthetic data collected under realistic experimental sampling conditions. (A) The accuracy of binarized promoter states *α*, averaged over 20 selected parameter sets (see main text for details). Comparisons are presented across three levels of transcriptional burstiness: high, intermediate, and low (from left to right). (B) An example trace highlighting DART’s improved accuracy over burstInfer in reconstructing the ground-truth binarized promoter states from the SSA for the case of low transcriptional bursting.

### 2.3 Statistical features from DART can distinguish models with different promoter states

Another key challenge is determining the number of underlying promoter states from live-cell imaging data, to select the most appropriate mechanistic model that accurately describes the observed dynamics. To accomplish this task, we used a machine learning-based model selection approach — a support vector-machine (SVM) (Materials and Methods sections 4.4; Fig. 1C).

To test the ability of SVM to distinguish between different models from ground-truth data, (i) the SVM was trained on several statistical features extracted from the distributions of the on- and off-times calculated directly from the SSA of a multi-state model of gene expression with one active state and a number of inactive states varying between 1 and 4 (illustrated in Fig. 6A); (ii) subsequently, new synthetic ground-truth distributions that it had not seen in its training were used to assess the classifier’s performance. Details can be found in Materials and Methods section 4.5.4. The results are summarized in Fig. 6B–D using confusion matrices, the Receiver Operating Characteristic (ROC) curve [51] and the Area Under the Curve (AUC) metric [51]. The values of the AUC are high, ranging between 0.81 and 0.92 (1 implying perfect distinguishability and 0.5 implying results are as good as picking a model randomly). The diagonal elements of the confusion matrices show that the percentage of correct classification varies from 64% for 2-vs 3-state models where the data is from the latter to 90% for 2-vs 4-state and 2-vs 5-state models where the data is from the 2-state model. This suggests that accuracy of model selection increases with the difference between the number of states of the true model and the proposed model, and that it is fundamentally difficult to distinguish between two models which are separated by only one state — further evidence is shown in Supplementary Fig. S5A. This intrinsic model indistinguishability stems from “model mimicry” whereby the statistical properties of a complex model are well approximated by simpler models [52].

**Fig. 6.**
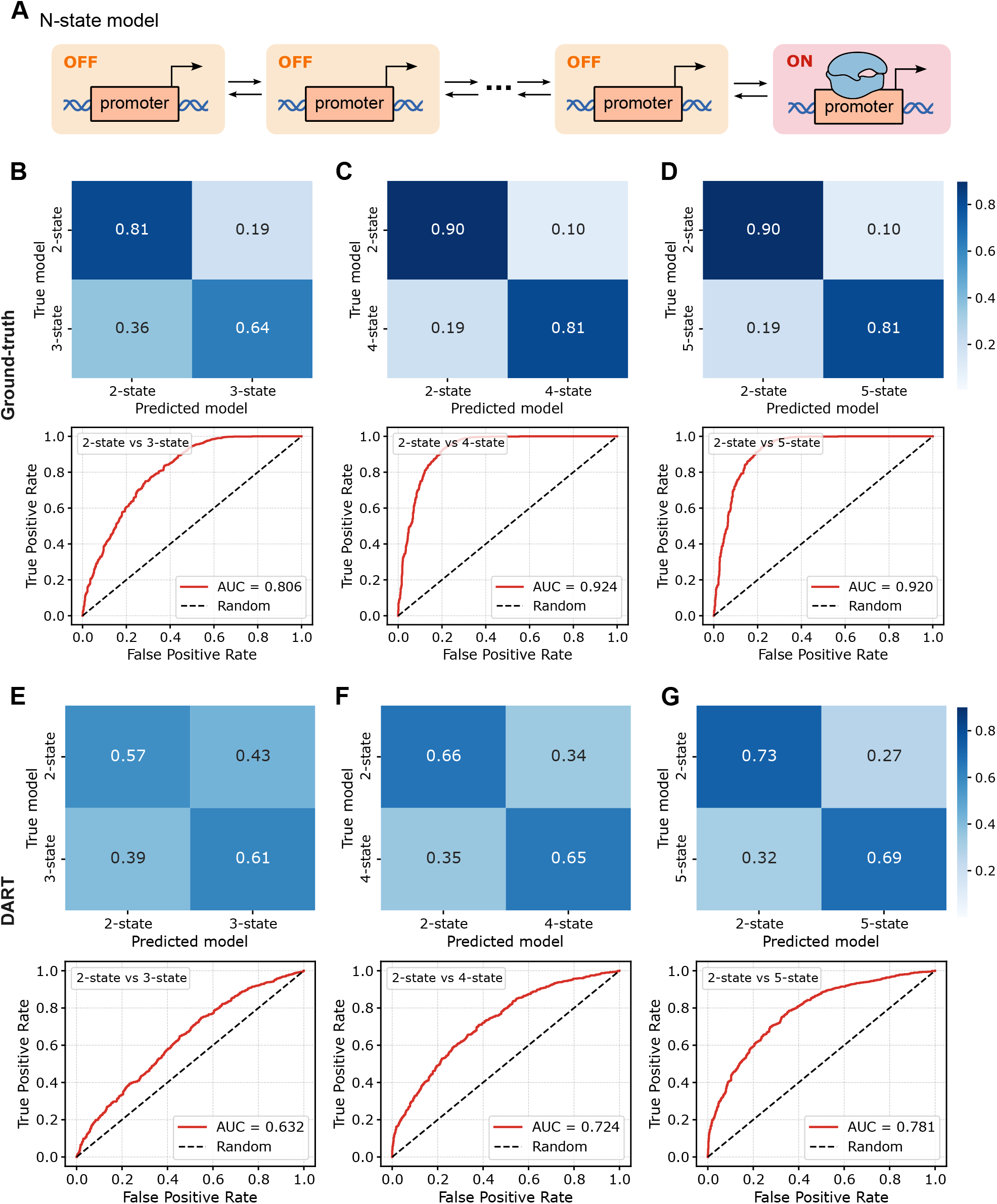
Model selection using a SVM classifier trained on the distributions of on- and off-times. (A) The classifier was trained and tested using synthetic data generated from an *N*-state model of stochastic gene expression with a single on-state and total number of states *N* varying between 2 and 5. (B-G) Classifier performance is evaluated through confusion matrices and ROC curves with corresponding AUC values, considering pairwise comparisons between the 2-state and 3-state models, the 2-state and 4-state models, and the 2-state and 5-state models. (B–D) Results correspond to SVM classification on features derived from the ground-truth distributions of on- and off-times from the SSA, whereas (E–G) show SVM classification on features extracted from the distributions of on- and off-times reconstructed by DART from SSA generated fluorescence data with 5% technical noise. The mean accuracy ⟨*α*⟩ of binarized promoter states are 0.79, 0.83, 0.85, 0.86 for the 2-, 3-, 4-, and 5-state models, respectively.

Next, we sought to understand the effect of DART’s binarization of fluorescence data on the SVM’s performance: (i) synthetic fluorescence data was generated from stochastic simulations of the multi-state model of gene expression shown in Fig. 6A with 5% lognormal distributed technical noise; (ii) the data was binarized by DART trained on a more general gene expression model comprising 3 on- and 3 off-states (as previously used for realistic experimental sampling protocols; for details see Materials and Methods, Section 4.5.3); (iii) statistical features were extracted from the distributions of on- and off-times output by DART and these were used to train the SVM to distinguish between models with varying numbers of states. Details can be found in Materials and Methods section 4.5.4. The results are summarized in Fig. 6E–G and Supplementary Fig. S5B. A comparison of the AUC and the confusion matrices with those of the SVM trained on ground-truth data (Fig. 6B–D and Supplementary Fig. S5A) shows that binarization has a significant effect on the SVM’s model selection performance, though the same trends that we previously found persist, i.e. model selection becomes more reliable when the proposed model and that underlying the data have large differences in the number of promoter states. Given that DART’s binarization accuracy is higher than that of common standard methods (Fig. 2), these results indicate that model selection using live-cell imaging data remains challenging and results should be used cautiously.

Similar issues have been reported using different approaches, such as BurstDECONV [42] which performs model selection using the reconstructed distribution of waiting times between successive polymerase initiation events from live-cell data and various other methods that distinguish models by means of the snapshot distributions of RNA counts [43, 53] — in fact, these apparently disparate approaches are very related because there is an exact relationship between the steady-state distribution of nascent RNA counts and the distribution of waiting times between successive polymerase initiation events [54].

### 2.4 DART reveals coupling between on-rates and off-rates in Drosophila even-skipped expression

Next, we consider the application of DART to study the transcriptional dynamics of the *even-skipped* (*eve*) gene in *Drosophila melanogaster* embryos, characterized by seven spatially periodic stripes during nuclear cycle 14 of the cellularizing blastoderm, immediately preceding gastrulation. Each of these stripes corresponds to a spatial domain along the anterior-posterior axis and is regulated by specific enhancers (top of Fig. 7A).

**Fig. 7.**
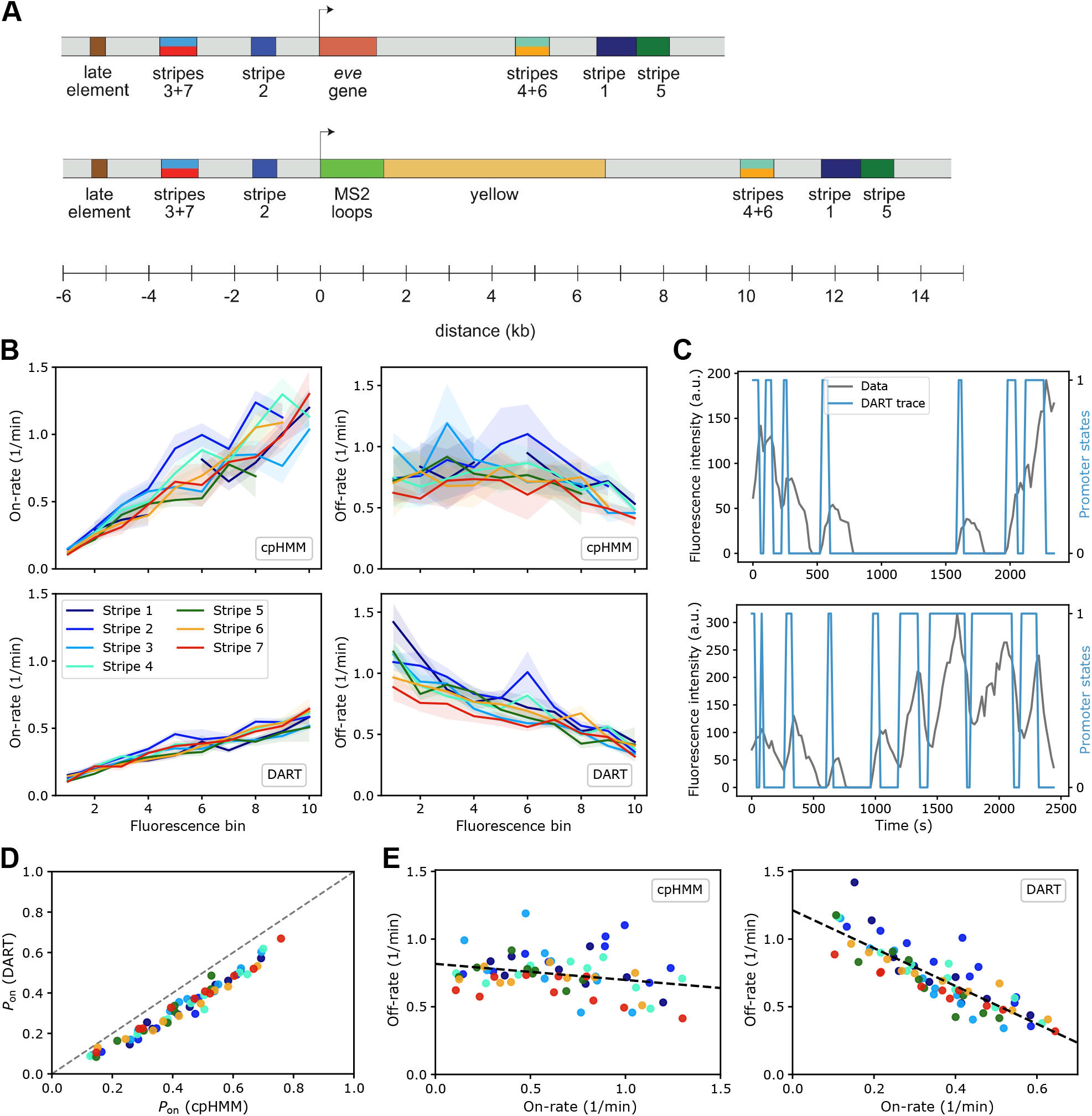
Application of DART to Drosophila *even-skipped* gene data **[55]**. (A) Top: Schematic of the wild-type *eve* gene locus including the five stripe enhancers and the late enhancer element. Bottom: Schematic of the engineered eve BAC, indicating the positions of the MS2 stem-loop array and the yellow reporter gene. Note these schematics were adapted from Fig. 1C and D of [55], licensed under CC-BY 4.0. (B) Comparison of inference results obtained with cpHMM (top) and DART (bottom). The left column shows inferred on-rates across 10 fluorescence intensity bins and 7 stripes, and the right column shows the corresponding inferred off-rates. Error bars represent standard deviations estimated by bootstrapping across multiple samples. (C) Representative examples of live-cell imaging data from [55], along with the corresponding promoter state trace inferred using DART. Examples are shown for particle ID 6.0124 (Stripe 6, fluorescence bin 2, top) and particle ID 1.0277 (Stripe 2, fluorescence bin 8, bottom). (D) Comparison of the inferred probability of the gene being in the on-state (*P*_on_) computed as the ratio on-rate/(on-rate + off-rate) from cpHMM (x-axis) and DART (y-axis) across all stripes and fluorescence intensity bins. Scatter points are colored according to stripe label. (E) Relationships between off-rate and on-rate inferred from cpHMM (left) and DART (right).

Berrocal et al. [55] used recombineering to modify a bacterial artificial chromosome (BAC) containing the *Drosophila melanogaster eve* locus, replacing the coding region with an MS2 stem-loop array followed by the yellow gene (for a schematic see the bottom of Fig. 7A). They observed that the nuclei in each stripe had a broad spread of average fluorescence values (implying a large heterogeneity in the number of RNAP molecules actively transcribing the gene). Because of the limited number of time points available in individual fluorescence traces, they grouped the nuclei within each stripe into ten fluorescence bins based on their mean fluorescence intensity. By binarization of the fluorescence traces using cpHMM [36], they calculated the rates at which the gene switches off and on as a function of the stripe label (1 *−* 7) and of the fluorescence bins in each stripe (1 *−* 10). The inferred rates, obtained from the file `w7_K3_t0_fluo_hmm_results_final.csv` provided by Berrocal et al. [55], are reproduced in Fig. 7B (top).

Next, we repeated the analysis but used DART instead of cpHMM. We first trained DART on synthetic data that was generated under the same experimental conditions employed in the Berrocal et al. paper [55] (Materials and Methods section 4.5.5). Subsequently, DART was applied to binarize the experimental fluorescence traces (obtained from the file `eve_data_longform_w_nuclei_060520.csv` in the Berrocal et al. paper). Two example traces (with stripe, fluorescence bin, and particle ID information from the same file) are shown in Fig. 7C. From the binarized time series, we calculated the rates at which the gene switches off and on as a function of the stripe label (1 *−* 7) and of the fluorescence bins in each stripe (for details of the calculations, see Materials and Methods Section 4.5.5). The results are shown in Fig. 7B (bottom). Comparing with Fig. 7B (top), we find that both cpHMM and DART predict that the switching on-rate increases with mean fluorescence (*R*^2^ = 0.77 and 0.83, respectively). In contrast, while cpHMM predicts an almost constant or a very weak decline of the switching off-rate with mean fluorescence, DART predicts a strong decline (*R*^2^ = 0.29 and 0.72, respectively). We note that cpHMM produced several NaN parameter estimates (6 out of 70 cases), resulting in discontinuities in the line plots of Fig. 7B(top). In all cases, DART yielded stable parameter estimates, underscoring the robustness of its switching rate estimation. Our analysis also shows that (i) cpHMM tends to sytematically underestimate the fraction of time spent in the on-state (Fig. 7D); (ii) cpHMM finds that the two switching rates are independent of each other (Fig. 7E left; *R*^2^ = 0.07) whereas DART finds a very clear linear-coupling between the two (Fig. 7E right; *R*^2^ = 0.73). Our findings are in agreement with those recently found by Chen et al. [56] in which a sophisticated Bayesian deconvolution framework was used to reconstruct transcription initiation events from live-imaging data.

## 3 Discussion

In this study, we introduced DART, a deep learning–based framework for the quantitative analysis of live-cell imaging data. We found that DART substantially improved the inference of promoter activity compared to conventional binarization methods. Across a wide range of biologically realistic synthetic datasets, DART consistently outperformed widely used approaches, including the two-state Hidden Markov Model and thresholding methods based on moving-average and Savitzky–Golay smoothing. Specifically, DART achieved *R*^2^ of 0.82 and 0.94 between predicted and true mean on- and off-times, respectively, whereas standard methods reached only 0.16 *−* 0.50. In addition, DART correctly classified promoter states in time with an accuracy exceeding 88%, even under conditions with strong memory effects in which fluorescence intensity at a given time point reflected previous rather than current promoter states. In such cases, standard methods not only exhibited reduced accuracy but also systematically overestimated the mean durations of the active and inactive states. DART also outperformed methods such as burstInfer, which were specifically designed to correct for memory in fluorescence signals. Notably, DART maintained high accuracy under realistic experimental constraints, including limited sampling rates and technical noise, highlighting its robustness and practical utility for the analysis of live-cell imaging data.

Beyond state inference, we demonstrated that statistical features derived from DART’s binarized traces can be used for gene expression model selection via a SVM classifier. We evaluated this approach in three scenarios in which ground-truth data were generated by a model with *M* states, and the SVM was tasked with distinguishing it from an alternative model with *N* states. Model selection accuracy was highest (80%) for discrimination between 2- and 4-state models (with ground truth from the 2-state model) and lowest (55%) for discrimination between 3- and 4-state models (with ground truth from the 4-state model). The reduced performance in certain cases reflected not a limitation of the DART–SVM framework itself, but rather the phenomenon of *model mimicry*, whereby the dynamics of a complex model can be closely approximated by simpler models. These findings suggest that binarized live-cell imaging data obtained under steady-state conditions are not optimally suited for reliably distinguishing between competing gene expression models. Instead, analyses based on time-dependent mRNA count distributions following perturbations, or on the snapshot joint distribution of nuclear and mature mRNA, may provide greater discriminatory power [18, 43, 44].

Finally, we applied DART to study the transcriptional dynamics of the *even-skipped* (*eve*) gene in *Drosophila melanogaster* embryos across different stripes and also within them. From DART’s binarized traces we computed the switching rates between the active and inactive states of the gene and found a strong linear coupling between the two rates. Notably, this coupling was not previously found when switching rates were estimated from binarization of the same data using cpHMM [55], a method that is more sophisticated than the bulk of commonly used methods because it corrects for memory in fluorescence signals. We note that the switching rate coupling that DART has elucidated is similar to that recently reported in [56].

An apparent current limitation of DART is its inability to predict the burst size as a function of time within live-cell imaging traces. However, this is easy to extract for a given binarized on-state identified by DART, by time-averaging the fluorescence intensity of the original signal over the on period. Further enhancements of DART’s accuracy for binarization of fluorescence time series with long-term memory effects could perhaps be obtained by using a different deep learning architecture based on transformers [57]. However, these improvements will likely only marginally improve DART’s predictions since it is already high. Another apparent limitation, is that we did not estimate mean on- and off-times for each individual cell, i.e. accounting for extrinsic noise. This was due to the nature of the fruit fly experimental data, where individual traces contained too few time points, rather than a limitation of DART itself. In fact, DART can perform cell-by-cell estimation provided that the fluorescence traces are sufficiently long.

Concluding, these results establish DART as a powerful and generalizable tool for quantitative analysis of transcriptional dynamics from live-cell imaging data. By leveraging deep learning, DART overcomes key limitations of traditional methods, enabling more accurate and robust inference of promoter state activity from single-cell data.

## 4 Materials and Methods

### 4.1 Commonly used baseline binarization methods

The three methods that we considered are: (1) A two-state hidden Markov model (HMM) was used to infer transcriptional states — each state corresponds to an on- or off-state. Initial states are determined by k-means clustering, and the model parameters are optimized through the Baum-Welch algorithm. The Viterbi algorithm is then employed to estimate the most likely sequence of hidden states, corresponding to the on and off states. The fast Julia implementation used in this paper is provided by HMMBase.jl [58]. We also obtained similar results using another implementation, HiddenMarkovModels.jl [59]; (2) A moving average smoothing method (MA) [34, 39] was initially applied using a convolution with a uniform kernel of 3 time points (i.e., a time-averaged window of three consecutive points). Following this, 10% of the maximum value of the smoothed fluorescence intensity data was set as the threshold. Fluorescence intensity values above this threshold were classified as on states, while values below were classified as off states; (3) The Savitzky-Golay (SG) filter [32] was applied to the fluorescence intensity data using a third-order polynomial and a window length of 11. The SG filter smooths data by fitting a low-degree polynomial to successive subsets of the signal via least squares regression, thereby preserving key features such as peak heights and widths while reducing noise. Following smoothing, a threshold corresponding to 10% of the maximum value of the smoothed fluorescence intensity data was determined and used to classify the promoter states as on or off.

### 4.2 Generation of synthetic data using a mathematical model of stochastic gene expression

#### 4.2.1 Stochastic Model

The model is defined by the reaction scheme:

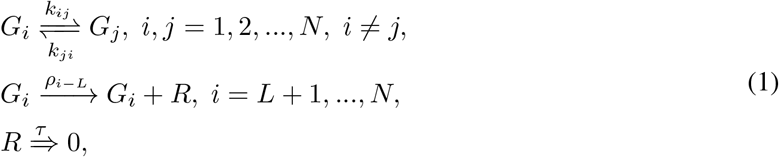

where *G*_*i*_ denotes the *i*-th gene state, with the first *L* states (*i* = 1, …, *L*) being off states, and the remaining *N − L* states (*i* = *L* + 1, …, *N*) being on states. Transitions from the *i*-th to the *j*-th gene state state occurs with a rate *k*_*ij*_. An RNA polymerase II (RNAP) *R* is produced from the *i*-th on-state at a rate *ρ*_*i−L*_. Each RNAP then undergoes elongation followed by termination which we assume to take a fixed duration *τ* — experimental support for the deterministic nature of the elongation and termination processes is provided in [60]. We note that this model is non-Markovian because the removal of *R* does not occur after an exponentially distributed time. Other studies have reported similar models of RNAP dynamics [54, 61, 62]. Simulations were performed using DelaySSAToolkit.jl [63], a Julia package designed for stochastic simulations with delays and implements several simulation methods proposed in [64, 65].

Next, we describe how, from the instantaneous RNAP dynamics, we calculate the total fluorescence intensity of nascent mRNA attached to RNAPs, which is the experimental observable. We will closely follow the approach described in [6]. In the stochastic simulations, each transcription initiation event produces an RNA polymerase II (RNAP) whose production time is recorded. Given the assumption of a constant elongation rate, the position of each RNAP at any subsequent time can be determined by multiplying the constant elongation rate by the time elapsed since its production. Once the position of the RNAP is known, the normalized fluorescence intensity *x*_*j*_ can be calculated using a simple function. This function is composed of two parts, a linear increasing part and a saturated constant part which model the underlying processes, as follows (for an illustration see Fig. 1A). As RNAP transcribes the engineered MS2 stem-loop sequences, which are specifically recognized by fluorescently tagged coat proteins, fluorescence intensity increases linearly due to the binding of fluorescent proteins to the nascent mRNA. The intensity associated with a single mRNA transcript reaches its maximum once the RNAP enters the gene body region. The simple function which quantifies this relationship is given by:

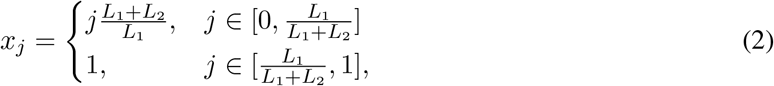

where *L*_1_ represents the length of the MS2 sequence, and *L*_2_ denotes the gene length. The variable *j* represents the normalized position of the RNAP on the gene. Thus, at any given time *t*, the total fluorescence intensity is obtained by summing *x*_*j*_ across all RNAP molecules present at that time.

#### 4.2.2 Selection of parameter values

For each parameter set, *N*_cell_ short stochastic simulation traces of duration *T* = 30 min were generated using the DelaySSAToolkit package, each representing an individual cell. These short traces were concatenated to form a single long trace with a total duration of *N*_cell_*T* which was subsequently sampled every Δ*t* time units (this will form the input to the deep learning model and the set of commonly used binarization methods). Furthermore, each concatenated trace was constrained to include at most one 30 minutes trace that did not generate any pulses (since such a trace provides no information on the switching dynamics between states). See later for a discussion about the values of *N*_cell_ and Δ*t*.

We define the mean duration of the total on-time (mean time spent in all on-states, ⟨*t*_on_⟩) and the mean duration of the total off-time (mean time spent in all off-states, ⟨*t*_off_⟩). Equations for ⟨*t*_on_⟩ and ⟨*t*_off_⟩ as a function of the gene expression model parameters are derived in the Supplementary Text S3. To cover the wide range of transcriptional burstiness seen *in vivo*, we selected parameter sets that generate three different levels of transcriptional burstiness. Gene expression appears bursty when the gene spends a short time in the on states (during which mRNA is produced in a burst) and a comparatively long time in the off states [66]. Hence, we chose the three levels of burstiness to be distinguished by the ratio of mean off-time to mean on-time ⟨*t*_off_⟩/⟨*t*_on_⟩: high burstiness: [5, 20], intermediate burstiness: [1, 5), and low burstiness [0.1, 1). Parameters were sampled using Sobol sequences [67], which are low-discrepancy, quasi-random sequences that cover the parameter space more uniformly than purely random sequences. Transition rates between states were drawn from the ranges inferred by fitting of the telegraph model to single-molecule Fluorescence In Situ Hybridization (smFISH) data from eukaryotic cells reported in the literature: *k*_*ij*_, *k*_*ji*_ *∈* [10^*−*3^, 10^1^] min^*−*1^, and synthesis rates from *ρ*_*i−L*_ *∈* [1, 50] min^*−*1^ [68, 69], subject to constraints on the burstiness (the ranges of ⟨*t*_off_⟩/⟨*t*_on_⟩ as discussed above) and the mean burst size *ρ*_bs_ *∈* [1, 150] [68]. Note that the mean burst size is the average amount of mRNA produced while the gene is in the on states (in the Supplementary Text S3, an expression is derived for the mean burst size in terms of the parameters of the model (1)). The time for an RNAP to elongate and terminate *τ* was restricted to [0.1, 10] min with the constraint that Λ = τ/⟨*t*_off_⟩ was in the range [0.1, 10] [70, 71].

A flowchart summarizing the generation of the parameter sets and the use of stochastic simulations to generate the synthetic fluorescence data is shown in Fig. 8A.

**Fig. 8.**
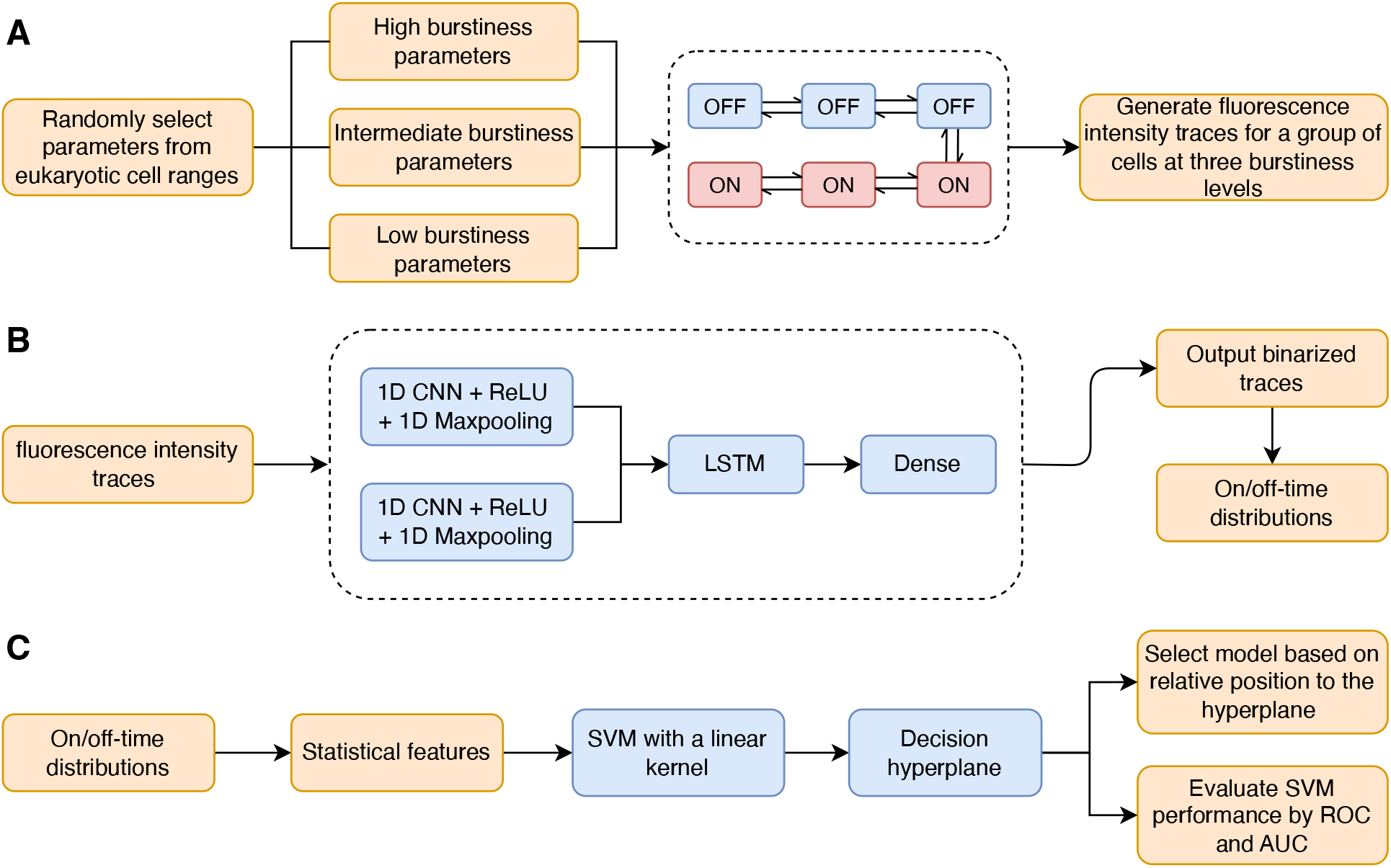
Flowchart summarising the training and evaluation of DART on synthetic data. (A) Synthetic fluorescence datasets were generated from stochastic simulations of a gene expression model with three off- and three on-states (1) to cover a broad range of transcriptional bursting scenarios (three levels ranging from low to high burstiness). (B) A deep learning architecture, comprising two CNN layers, one LSTM layer, and one dense layer, was trained on the synthetic datasets from each burstiness level. The deep learning model outputs binarized state traces, where a value of 1 denotes an on-state and 0 denotes an off-state. From these traces, on- and off-time distributions were extracted. (C) Statistical features derived from these distributions were used as input for training a SVM classifier. The trained SVM model performs model selection by classifying new test samples based on their relative position to the decision hyperplane. The classifier’s performance of distinguishing between models was assessed using receiver operating characteristic (ROC) curves and the area under the curve (AUC) metric.

### 4.3 Deep learning architecture for trace binarization

The concatenated, long traces of synthetic fluorescence intensity data were input to a deep learning model developed to binarize the trace, i.e. to infer gene on- and off-states. The deep learning architecture consists of 2 one-dimensional convolutional neural network (1D-CNN) layers [72], followed by a Long Short-Term Memory (LSTM) layer [73, 74], and a dense output layer.

To extract local temporal features from the fluorescence signals, we employed the 1D-CNN as the initial processing stage. Each convolutional layer applies a set of learnable kernels ***w*** ∈ ℝ^*l*^ across the input sequence 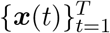, producing feature maps that capture patterns within fixed-size temporal windows. Following convolution, a Rectified Linear Unit (ReLU) activation function [75] is applied to introduce non-linearity and improve learning capacity. We further perform 1D max pooling with a fixed window size and stride to reduce the temporal dimension and retain dominant local features. To preserve the input sequence length and avoid boundary effects, we apply symmetric zero-padding during both convolution and pooling operations. The sequence 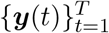, obtained by passing 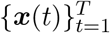 through two successive convolutional layers, is used as the input to a LSTM layer [73, 74].

LSTMs are a specialized class of recurrent neural networks (RNNs)[74, 76] designed to address the vanishing gradient problem [77], which limits the ability of standard RNNs to capture long-term dependencies in sequential data. LSTMs use a memory cell and gating mechanisms to selectively retain, update, and output information over time, enabling the modeling of long-term dependencies in sequential data. In our setting, the LSTM output at each time step is passed to a dense layer with a sigmoid activation function to produce a binary output 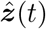, representing the inferred on/off promoter state underlying the original fluorescence signal ***x***(*t*), where 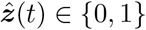, with 1 denoting the on state and 0 indicating the off state.

So in summary, the convolutional layers are employed to capture local temporal features and correlations along the input signal, while the LSTM layer captures longer-range dependencies across the time series.

The whole deep learning model is trained end-to-end using backpropagation to minimize the binary cross-entropy loss:

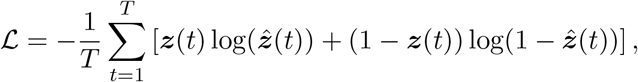

where ***z***(*t*) is the true gene state’s trace (with each element of ***z***(*t*) belonging to the set *{*0, 1*}*) generated from the SSA, and 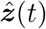 is the deep learning model’s prediction. On- and off-time distributions can then be calculated from the predicted binary traces. All implementations of deep learning were carried out using Flux.jl [78], a Julia [79] package specifically designed for deep learning applications. For details of hyperparameter tuning, see Supplementary Text S4 and Supplementary Fig. S6. A flowchart summarizing the deep learning model architecture is shown in Fig. 8B.

### 4.4 SVM classifier for model selection

To perform model selection among promoter switching models with different numbers of on- and off-states, we employed a Support Vector Machine (SVM) classifier with a linear kernel [80, 81]. SVM is a supervised machine learning method widely used for classification tasks. It classifies data by identifying an optimal decision boundary (hyperplane) that separates data points belonging to different classes. Given that our task involves classification among multiple promoter switching models, we implemented a one-vs-one (OvO) classification strategy, training an SVM classifier for each pair of models. For a classifier distinguishing between models *i* and *j*, parameter sets corresponding to model *i* were labeled positively (*m*_*i*_ = +1), while those from model *j* were labeled negatively (*m*_*j*_ = *−*1); samples from all other models were excluded from training.

The distributions of on- and off-times obtained from the ground-truth promoter state data (SSA) or from binarized traces produced by the deep learning model were used. To characterize the statistical properties of these distributions, we extracted a set of statistical features ***v***_*i*_, where *i* = 1, 2, …, *S*, and *S* denotes the total number of parameter sets. These features included the mean, standard deviation, coefficient of variation (CV), the maximum values of the on- and off-times, the proportion of on- and off-times exceeding specific thresholds and three different percentiles (*Q*_1_ = 0.25, *Q*_2_ = 0.50, *Q*_3_ = 0.75). The resulting feature vectors were subsequently used as inputs to train the SVM classifier (For training details, see Section 4.5.4). The classifier’s performance was assessed by the Receiver Operating Characteristic (ROC) curve [51] and the Area Under the Curve (AUC) metric [51]. For each test parameter set, the promoter model corresponding to the highest decision value was selected, with higher predicted probabilities indicating greater classification confidence. The SVM classifier in this paper was implemented using the Scikit-learn library in Python [82]. For details of hyperparameter tuning, see Supplementary Text S4. A flowchart summarizing the use of the SVM for model selection is shown in Fig. 8C.

### 4.5 Synthetic data generation and training process

#### 4.5.1 Synthetic data generated with optimized sampling to evaluate baseline method accuracy

The long concatenated traces of simulated fluorescence intensity (discussed in Section 4.2.2) were used to test the accuracy of the three baseline methods in Section 4.1. Specifically, the period Δ*t* between successive measurements of the intensity in a trace was chosen as Δ*t* = 0.1 *×* min(⟨*t*_on_⟩, ⟨*t*_off_⟩) and the cell number *N*_cell_ is selected as the minimum of 1000 and the value of *N*_cell_ in Eq. (3) that corresponds to *N*_pulse_ = 1100.

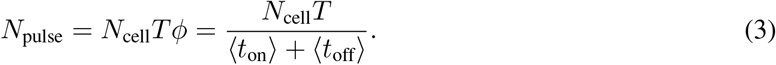

Note that *N*_pulse_ is the average number of pulses corresponding to transcriptional on-times in the long trace, *ϕ* = 1/(⟨*t*_on_⟩ + ⟨*t*_off_⟩) is the pulse frequency. The upperbound constraint of 1000 is chosen to mirror the fact that live-cell imaging experiments typically report data for less than a thousand cells. This setting is an idealized scenario where the sampling interval and the total cell number are chosen such that the long traces always have sufficient statistical information about the underlying promoter switching dynamics, i.e. burst events can always be detected for any parameter set. This procedure led to the following ranges: Δ*t ∈* [0.21, 26.4] seconds and *N*_cell_ *∈* [4, 555].

In addition, in this idealized setting, we assumed that all MS2 sequence loops bind to the fluorescence probe instantaneously upon transcription, resulting in an immediate rise in fluorescence intensity to its maximum level, without a gradual linear increase, i.e. the ratio *L*_1_*/*(*L*_1_ + *L*_2_) = 0 in Eq. (2) and Fig. 1A. The synthetic data was generated under these conditions from stochastic simulations of the model in Eq. (1) with *L* = 3, *N* = 6 (corresponding to 3 off- and 3 on-states) for three groups of 600 parameter sets that lead to high, intermediate, and low levels of transcriptional burstiness (a total of 1800 parameter sets).

#### 4.5.2 Training the deep learning model on optimally sampled synthetic data

The deep learning model was trained on simulated data generated under the same idealized conditions, i.e. *L, N*, Δ*t, N*_cell_ and *L*_1_ were chosen as in Section 4.5.1. The data was obtained by simulations using three groups of parameters sets covering the three burstiness levels. Each group consisted of 3300 parameter sets, with 2400 used for training, 300 for validation, and 600 for testing the deep learning model. Note that the latter 600 parameters are the same as the 600 parameters (for each burstiness level) used to test the accuracy of the baseline methods in Section 4.5.1.

#### 4.5.3 Training the deep learning model on realistically sampled synthetic data

A different setting was used to generate synthetic data under realistic experimental sampling protocols, where Δ*t* was fixed to 6 seconds and the cell number *N*_cell_ to 300. This reflects a typical experimental scenario in which the sampling interval and the total cell number are fixed. In this setting, for simplicity, the ratio *L*_1_*/*(*L*_1_ + *L*_2_) was set to zero.

The deep learning model was then trained on synthetic fluorescence traces simulated from the the model described in Eq. (1), with parameters *L* = 3 and *N* = 6. To account for experimental background noise, we added log-normal noise with the same mean as that of the fluorescence traces and with coefficients of variation (CV) of 5%, 10%, 20%, and 30%. Three groups of parameter sets were considered, corresponding to high, intermediate, and low burstiness levels (Section 4.2.2). In addition, to reflect the experimental assumption that transcriptional bursts can be reliably detected at the temporal resolution of 6s, we restricted parameter sets to cases where the mean on-time was greater than or equal to the sampling interval. Each burstiness level group contained 3700 parameter sets, where 2400, 300, and 1000 parameter sets were used for training, validation, and testing of the deep learning model, respectively.

#### 4.5.4 SVM for realistically sampled synthetic data

Stochastic models used for model selection included the N-state stochastic model with one on-state and one, two, three, or four off-states (corresponding to Eq. (1) with *L* = 1, 2, 3, 4 and *N* = 2, 3, 4, 5, respectively). For each model of gene expression, 4000 parameter sets were randomly sampled using Sobol sequences within biologically realistic ranges for eukaryotic cells exhibiting an intermediate level of transcriptional burstiness level (Section 4.2.2). Consistent with the setup in Section 4.5.3, the sampling interval was set to Δ*t* = 6 seconds, the number of cells to *N*_cell_ = 300, and the ratio *L*_1_*/*(*L*_1_ + *L*_2_) was set to 0.

Statistical feature vectors were extracted from the distributions of on- and off-times (see Section 4.4). These feature vectors were then used as inputs to train the SVM classifier to distinguish between stochastic models with one on-state and one, two, three or four off-states. The dataset was partitioned into a training set (80%) and a test set (20%). 5-fold cross-validation was then applied to the training set to ensure robustness.

#### 4.5.5 Deep learning for *Drosophila* data

The stochastic model in Eq. (1), with parameters *L* = 3 and *N* = 6 was used to generate synthetic data to train the deep learning model. 3700 parameter sets were randomly sampled using Sobol sequences within biologically realistic ranges for eukaryotic cells (Section 4.2.2) with the difference that ⟨*t*_off_⟩/⟨*t*_on_⟩ was sampled from the broad range [1, 15] corresponding to both intermediate and high burstiness levels of gene expression. Consistent with the experimental setup in Berrocal et al. [55], the sampling interval was set to Δ*t* = 20 seconds, *L*_1_ *≈* 1500 base pairs (MS2 sequence length), *L*_2_ *≈* 5165 base pairs (yellow gene length) and an elongation time *τ ≈* 2.33 minutes. Due to the limited number of time points available in individual fluorescence traces, the original study grouped nuclei within each stripe into ten fluorescence bins based on their mean fluorescence intensity to facilitate inference. Therefore, the number of cells was set to *N*_cell_ = 40 in our simulation, corresponding to the approximate number of cells in each fluorescence bin group. In addition, 5% log-normal distributed technical noise was added. Out of the total of 3700 parameter sets, synthetic traces from 2400 of these were used to train the deep learning model, and 300 and 1000 were used for validation and testing, respectively.

Subsequently, the experimental fluorescent time-series data from Berrocal et al. [55] was input to the deep learning model to infer the underlying time series of binarized promoter states, and the effective on- and off-rates were estimated as

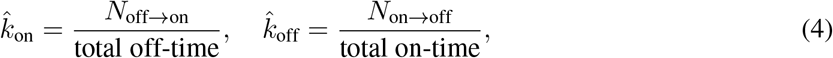

where *N*_off*→*on_ denotes the number of transitions from the off-state to the on-state, and *N*_on*→*off_ denotes the number of transitions from the on-state to the off-state. These definitions are intuitive and do not make strong assumptions about the nature of the distributions of the on- and off-times. The on- and off-rates were separately computed for each fluorescence bin group using data pooled across all cells in that group to ensure sufficient sampling.

## Data availability

The code for this paper is available at https://github.com/mmmuhan/DART.

## Acknowledgments

M. M. acknowledges support from a PhD scholarship provided by the Darwin Trust. R. G. acknowledges support from the Leverhulme Trust (RPG-2024-082). The authors thank Jun Liu for his helpful advice on machine learning and Augustinas Sukys for his valuable feedback on previous drafts of this work.

## Supplementary Text S1: Application of DART to synthetic data generated by simulation of a different class of gene expression models

Another widely studied class of promoter switching models is one in which gene states are connected by effective irreversible reactions [40, 43, 83]. Specifically, in this model a gene transitions sequentially through *N −* 1 inactive states before reaching a single active state *G*_*N*_. Upon exiting the active state, the promoter returns directly to the initial inactive state, forming a closed-loop transition cycle:

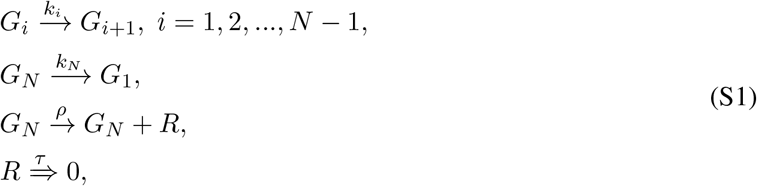

where *G*_1_, …, *G*_*N−*1_ are off states, and *G*_*N*_ is the single on-state. Transcription occurs in the on-state at a rate *ρ*, followed by transcript elongation and detachment over a fixed duration *τ*. Note that because of the reaction *G*_*N*_ *→ G*_1_, this model is not a special case of the reversible model of gene expression (1). From the instantaneous RNAP dynamics of an stochastic simulation of this model, one can calculate the total fluorescence intensity of nascent mRNA attached to RNAPs using the algorithm described in Materials and Methods Section 4.2.1.

The trained deep learning model from Section 4.5.2 was applied to evaluate the generalizability of our approach, i.e. to test the ability of DART to binarize synthetic data generated using a different model than it was originally trained on. The test data set consisted of idealized synthetic data generated from the three-state irreversible model in Eq. (S1), with *N* = 3 and Δ*t, N*_cell_ and *L*_1_ chosen as in Section 4.5.1. A total of 600 parameter sets were randomly sampled from eukaryotic cells ranges using Sobol sampling (see Section 4.2.2 for details on parameter selection), at a high burstiness level as a representative case. As shown in Fig. S1, DART accurately inferred both on- and off-times, despite the altered structure of the irreversible model compared to the reversible model used for training.

## Supplementary Text S2: Robustness of DART on idealized data

To assess the robustness of the deep learning model architecture implemented in DART, we performed 100 independent training and evaluation runs on the same optimally sampled synthetic data (Section 4.5.2 in Materials and Methods), each initialized with a different random seed controlling parameter initialization and data shuffling during training. This approach enables quantification of variability in model performance attributable solely to variations introduced during the training process, such as weight initialization and batch sampling. For each of the 100 runs, we computed the three evaluation metrics described in the main text: (1) mean binary vector accuracy ⟨*α*⟩, (2) mean relative error (MRE) of the estimated mean on-time, and (3) MRE of the estimated mean off-time. Each metric value corresponds to the average performance over a test set of 600 parameter sets for one of 100 random seeds. The resulting values were then aggregated across all runs to construct posterior distributions, which are shown in Fig. S2. The performance is further evaluated across the three burstiness levels defined in Materials and Methods Section 4.2.2. The results show that the deep learning model implemented in DART maintains consistent performance across all runs with different random seeds, demonstrating robustness to inherent variability introduced during the training process.

## Supplementary Text S3: Useful stochastic properties of the gene expression model

Here, we derive analytical expressions for the mean on-time, the mean off-time, and the mean burst size for the gene expression model (Eq. (1)) with 3 on- and 3 off-states, i.e. *L* = 3, *N* = 6. The reaction scheme is

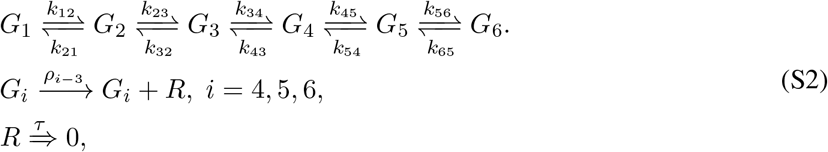

where the first three promoter states (*G*_1_-*G*_3_) are off, and the remaining three (*G*_4_-*G*_6_) are on.

### Calculation of the mean off-time

The off-time starts when the promoter switches from state *G*_4_ to state *G*_3_ and ends when it switches from *G*_3_ back to *G*_4_. The distribution of the off-time can be formulated as a first-passage time problem [84]. Specifically, we consider the sub-network

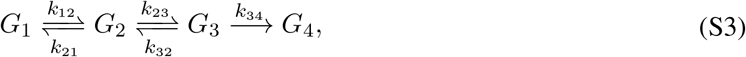

with the initial condition that at *t* = 0, the system is in state *G*_3_; our aim is to find the mean of the first-passage time distribution to exit to *G*_4_.

Let *P*_*i*_(*t*) denote the probability that the promoter is in state *G*_*i*_ at time *t* given that it was in state *G*_3_ at *t* = 0. The stochastic dynamics are described by the following set of differential equations:

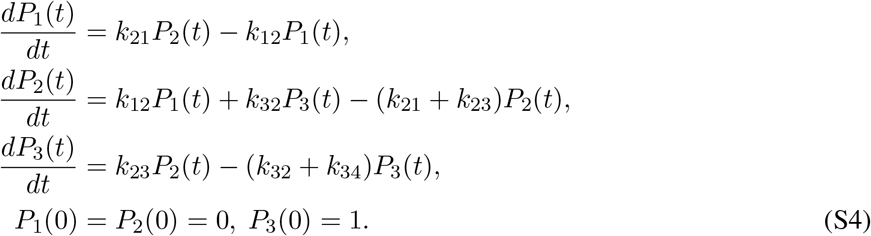

Taking the Laplace transform, we obtain

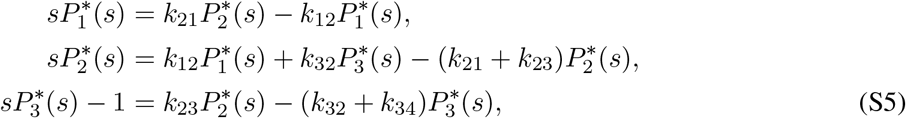

where 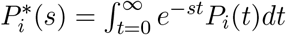. This system of equations can be solved simultaneously to obtain 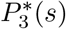. The distribution of the off-time is then given by *f* (*t*) = *k*_34_*P*_3_(*t*) and hence the mean off-time is given by

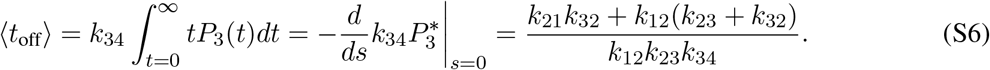

### Calculation of the mean on-time

The on-time starts when the promoter switches from state *G*_3_ to state *G*_4_ and ends when it switches from *G*_4_ back to *G*_3_. For this calculation we only need to consider the sub-network

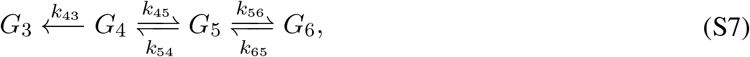

with the initial condition that at *t* = 0, the system is in state *G*_4_. The aim is to find the mean of the first-passage time distribution to exit to off-state *G*_3_.

Let *P*_*i*_(*t*) denote the probability that the promoter is in state *G*_*i*_ at time *t* given that it was in state *G*_4_ at *t* = 0. The stochastic dynamics are described by the following set of differential equations:

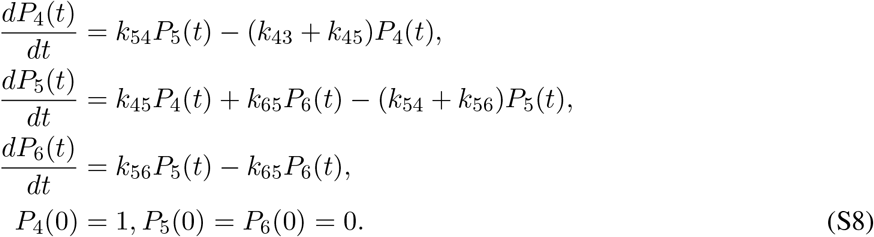

As we did for the mean off-time calculation, we can obtain the mean on-time via the Laplace transform method:

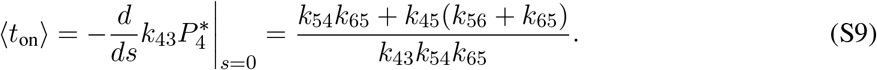

### Calculation of the mean burst size

The mean burst size is the total mean mRNA produced while the system is in one of its 3 on-states. Note that this definition does not account for any mRNA degradation.

The system of differential equations consists of Eq. (S8) and an additional equation

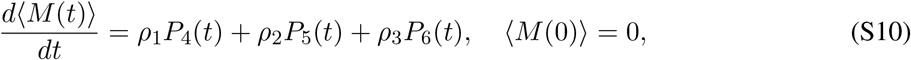

where ⟨*M* (*t*)⟩ is the mean mRNA produced at time *t*. This system of 4 differential equations can be solved using the Laplace transform method. The mean burst size is then given by

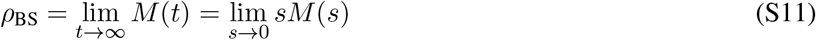

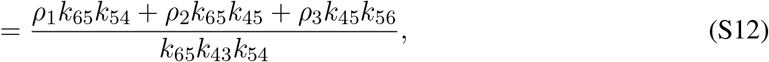

where Eq. (S11) follows from the final value theorem. Note that the intuition behind this calculation is that the amount of mRNA will increase monotonically with time while the promoter is active and will saturate to a constant (*ρ*_BS_) when the promoter switches to an inactive state. Note also that for the special case of models with only one active state, such as the conventional telegraph model, we have *k*_45_ = 0, which recovers the well-known result that the mean burst size is equal to *ρ*_1_*/k*_43_.

## Supplementary Text S4: Hyperparameter Tuning

Hyperparameter tuning was performed during training to optimize the deep learning model’s and SVM classifier’s performance and generalization. Here, we discuss the choice of batch size, learning rate, number of convolutional layers, kernel size of the convolutional layer, number of filters per layer and values of the regularization parameters.

Our deep learning framework was trained using the Adam optimizer [85], which adapts learning rates for individual parameters based on first- and second-moment estimates of past gradients. Network weights were initialized using the Glorot Uniform initialization method [86], which helps maintain stable gradients at the start of training.

The batch size influences both training dynamics and generalization. Smaller batch sizes introduce more stochasticity into the gradient estimates, which can help the model escape sharp minima and often improve generalization, but at the cost of slower convergence. Larger batch sizes yield more stable and efficient gradient estimates, leading to faster training, but they can bias optimization toward sharper minima, which are sometimes associated with poorer generalization. In this study, we set the batch size to 20, as it consistently yielded good performance across numerical experiments.

The learning rate is another key hyperparameter in the training process as it controls the step size of weight updates and thus influences the convergence behavior. We chose a learning rate scheduling strategy similar to that proposed in [87]. The initial learning rate was set to 0.01 and was halved if the average validation loss over the previous 25 epochs improved by less than 0.005. The training was terminated when the learning rate had been reduced six times or when the number of epochs reached 200. We found that this approach ensured the convergence of the loss function across our experiments.

We systematically explored different combinations of these hyperparameters and and identified two candidate neural network architectures. The first architecture, denoted as 3-3, comprises two convolutional layers, each with a kernel size of 3, followed by a single LSTM layer and a final dense layer, with successive feature dimensions of 32, 64, and 64, respectively. The second architecture, denoted as 3-3-3, consists of three convolutional layers, each with a kernel size of 3, followed by an LSTM layer and a dense layer, with feature dimensions of 32, 64, 64, and 64, respectively. As shown in Fig. S6, three metrics were used to evaluate and compare the performance of the two candidate architectures: mean binary vector accuracy ⟨*α*⟩, and the MRE of the estimated mean on- and off-times. As in Supplementary Text S2, each metric was computed as the average performance over a test set of 600 parameter sets for a given random seed (out of 50 in total). The posterior distribution for each metric was then constructed across these 50 seeds. The results indicate that the two architectures performed similarly for binary vector accuracy but the 3-3 architecture consistently outperformed the alternative across all three burstiness levels using the MRE metric. Therefore, the 3-3 structure was selected as the optimal architecture for DART.

In addition to the deep learning model, we also optimized the key hyperparameter of the SVM classifier. A grid search was performed to identify the optimal value of the regularization parameter *C* that controls the trade-off between maximizing the margin and minimizing classification errors. Values of *C* were sampled uniformly in logarithmic space over the interval [0.001, 1000], yielding 100 candidate values. Performance was evaluated based on the average one-vs-one AUC score across all model classes. The optimal value identified through this procedure was *C* = 0.02.

## Supplementary Figures

**Fig. S1.**
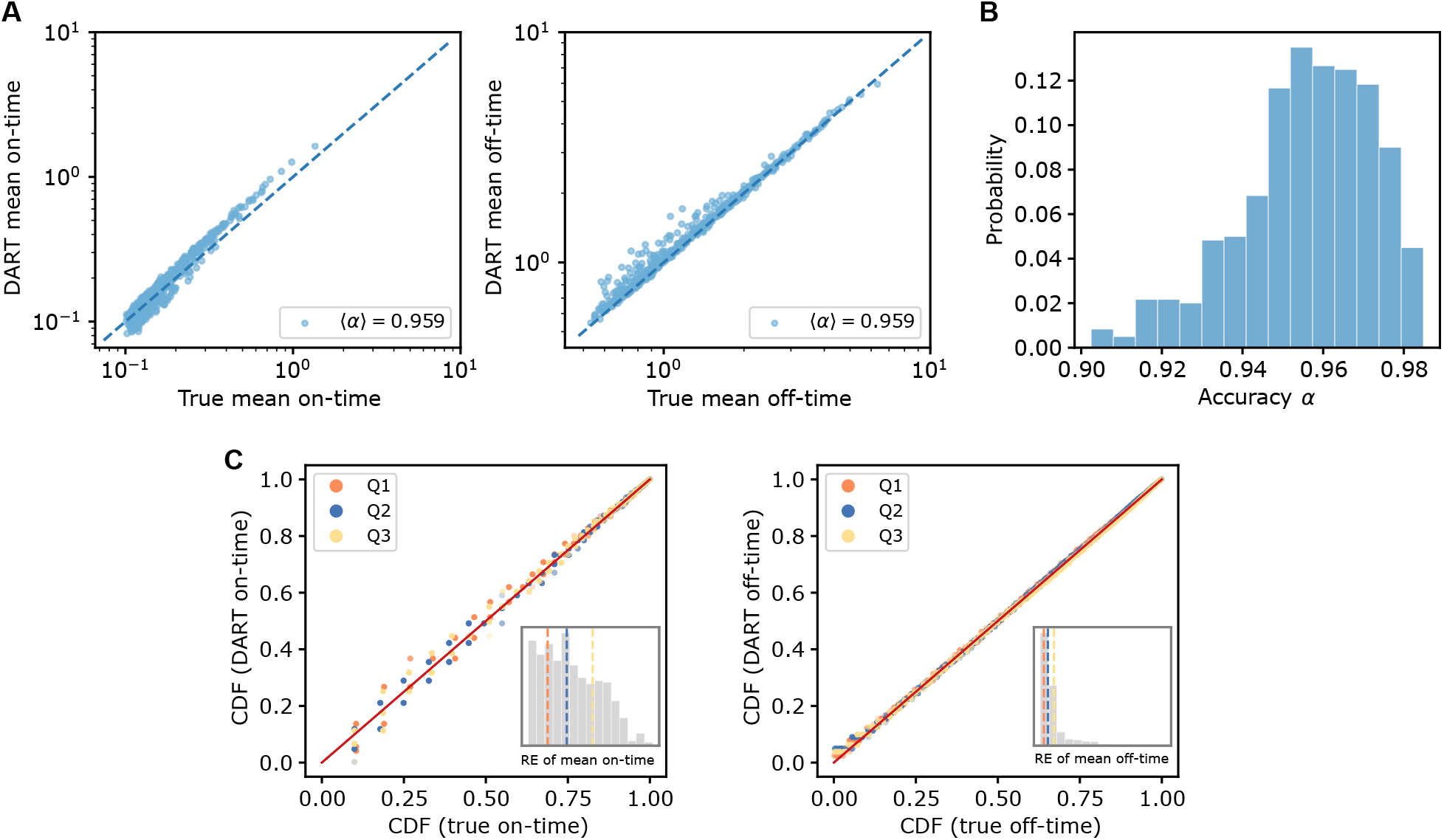
Binarization performance of DART on synthetic data generated by a model of gene expression — 3-state irreversible model with a high level of transcriptional bursting — that it has not seen in its training. (A) Comparison between the true mean on- and off-times obtained from the SSA and those inferred by DART. (B) Distribution of the proportion of correctly classified promoter states in time (the binary state vector accuracy *α*) produced by DART. (C) P–P plots comparing the CDFs of true on/off-times generated by the SSA with the CDFs of the inferred on/off-times from DART. Parameter sets corresponding to the Q1, Q2, and Q3 quartiles of the RE distribution of the mean on- and off-times were selected for illustration. See Supplementary Text S1 for details.

**Fig. S2.**
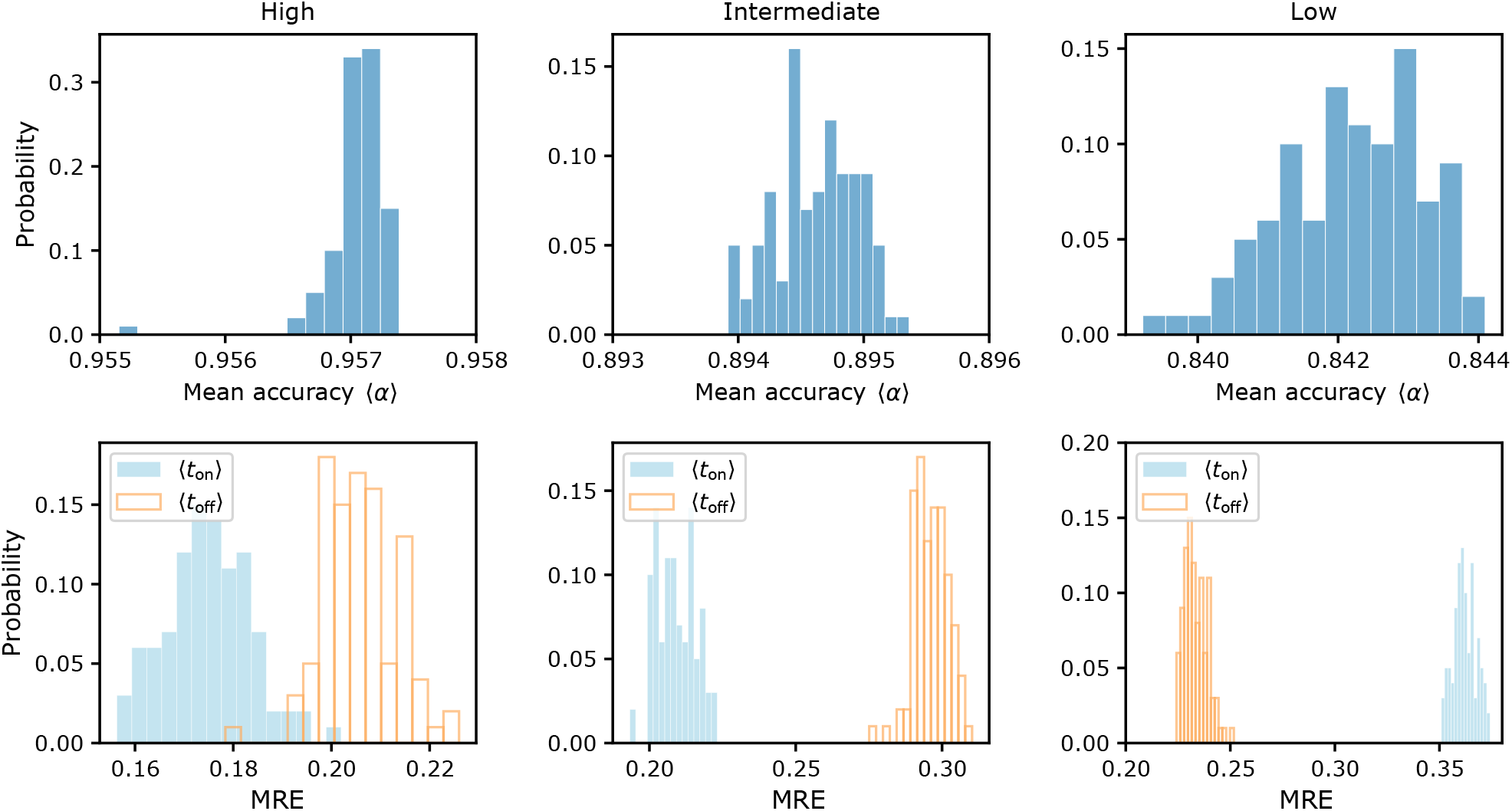
Posterior distributions of the binary vector accuracy *α* and the MRE over 100 random seeds for DART. Each column represents a different burstiness level: left—high, middle—intermediate, and right—low. The first row shows the posterior distribution of the mean binary vector accuracy ⟨*α*⟩, while the second row shows the posterior of the MRE of the estimated mean on- and off-times. DART was trained using 100 random seeds on identical idealized synthetic data (see Supplementary Text S2 for details).

**Fig. S3.**
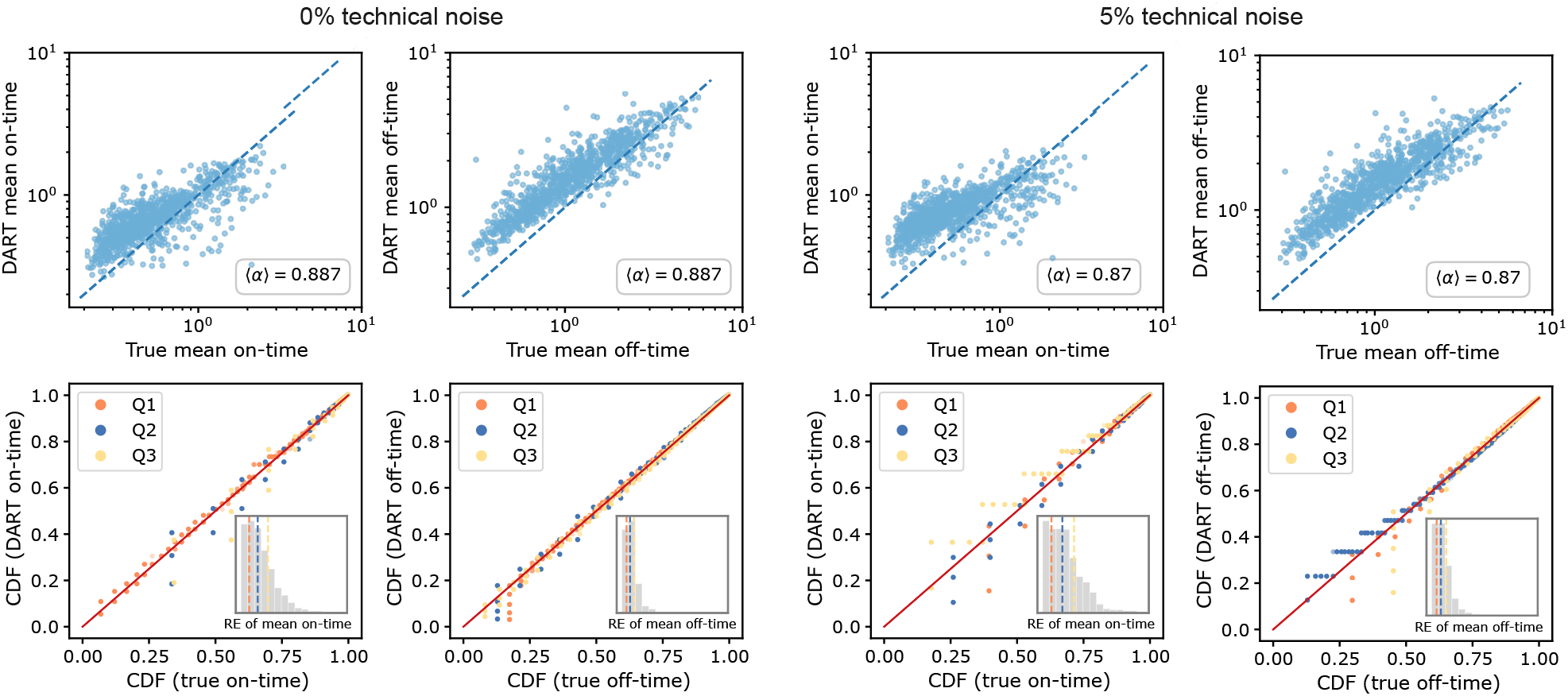
Performance of DART using synthetic data collected under realistic experimental sampling protocols and an intermediate level of transcriptional bursting. The first row compares the true mean on- and off-times from the SSA with the estimates from DART. The second row presents P–P plots comparing the true CDFs generated via the SSA with those produced by DART, evaluated at the 0.25 (Q1), 0.5 (Q2), and 0.75 (Q3) quantiles of the RE distribution of the mean on- and off-time estimates. The left panel represents the case without technical noise, while the right panel shows the case with 5% technical noise (log-normal distributed). For more details, see Main text section 2.2.

**Fig. S4.**
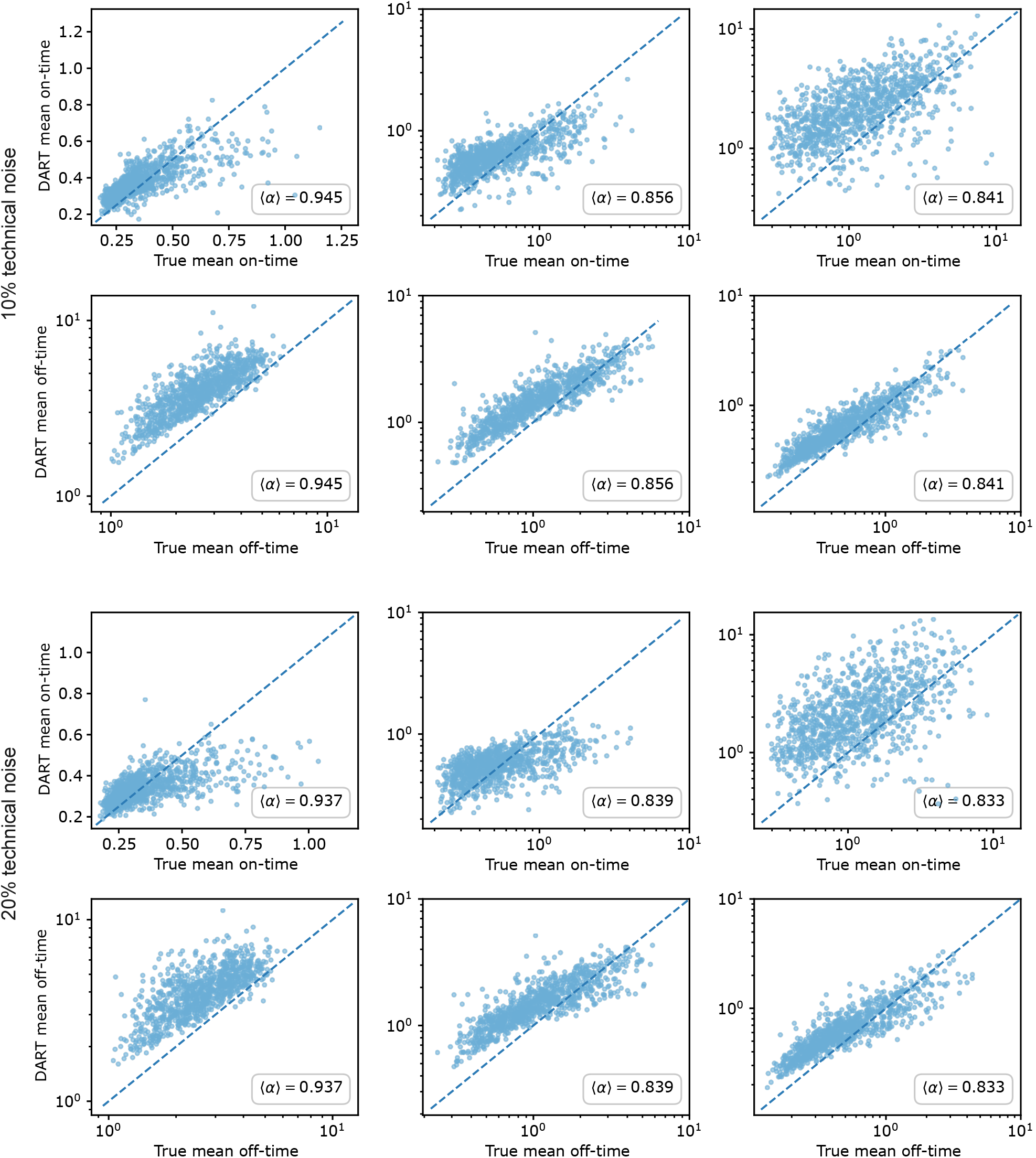
Performance of DART under realistic experimental sampling protocols with varying levels of technical noise. The top panel corresponds to log-normal distributed technical noise with a coefficient of variation (CV) of 10%, while the bottom panel shows results for noise with a CV of 20%. In each panel, the first row shows the model’s performance in estimating the mean on-time relative to the ground true values from the SSA, while the second row shows the performance in estimating the mean off-time. DART was trained separately on realistic synthetic data with 10% and 20% technical noise, and its performance was evaluated on a test set of size 1000, following the procedure described in Materials and Methods, Section 4.5.3.

**Fig. S5.**
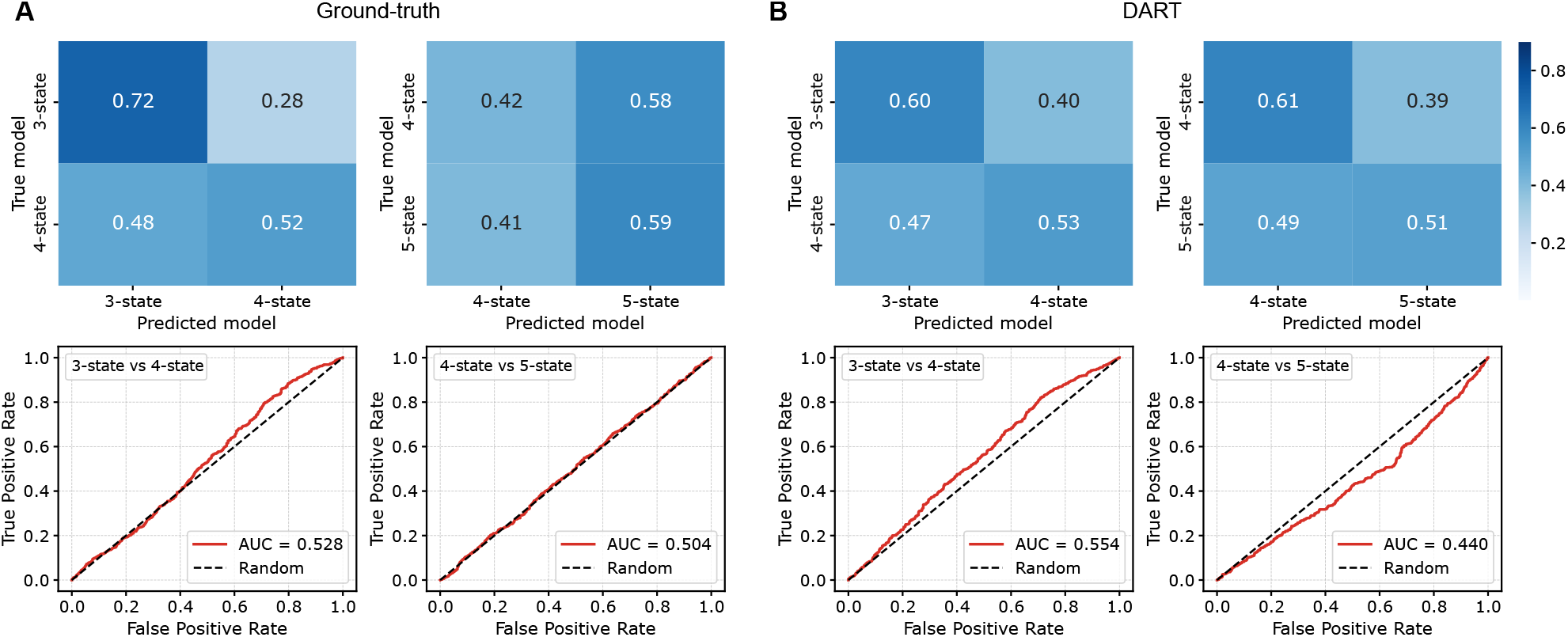
SVM-based model selection using the the distributions of on- and off-times. (A) Classification based on features of the ground-truth distributions of the on- and off-times generated from the SSA. (B) Classification based on distribution features extracted by DART from synthetic fluorescence data generated by the SSA and with 5% technical noise. Details as in Fig. 6, except that model selection here is exclusively between 3- and 4-state models, and between 4- and 5-state models.

**Fig. S6.**
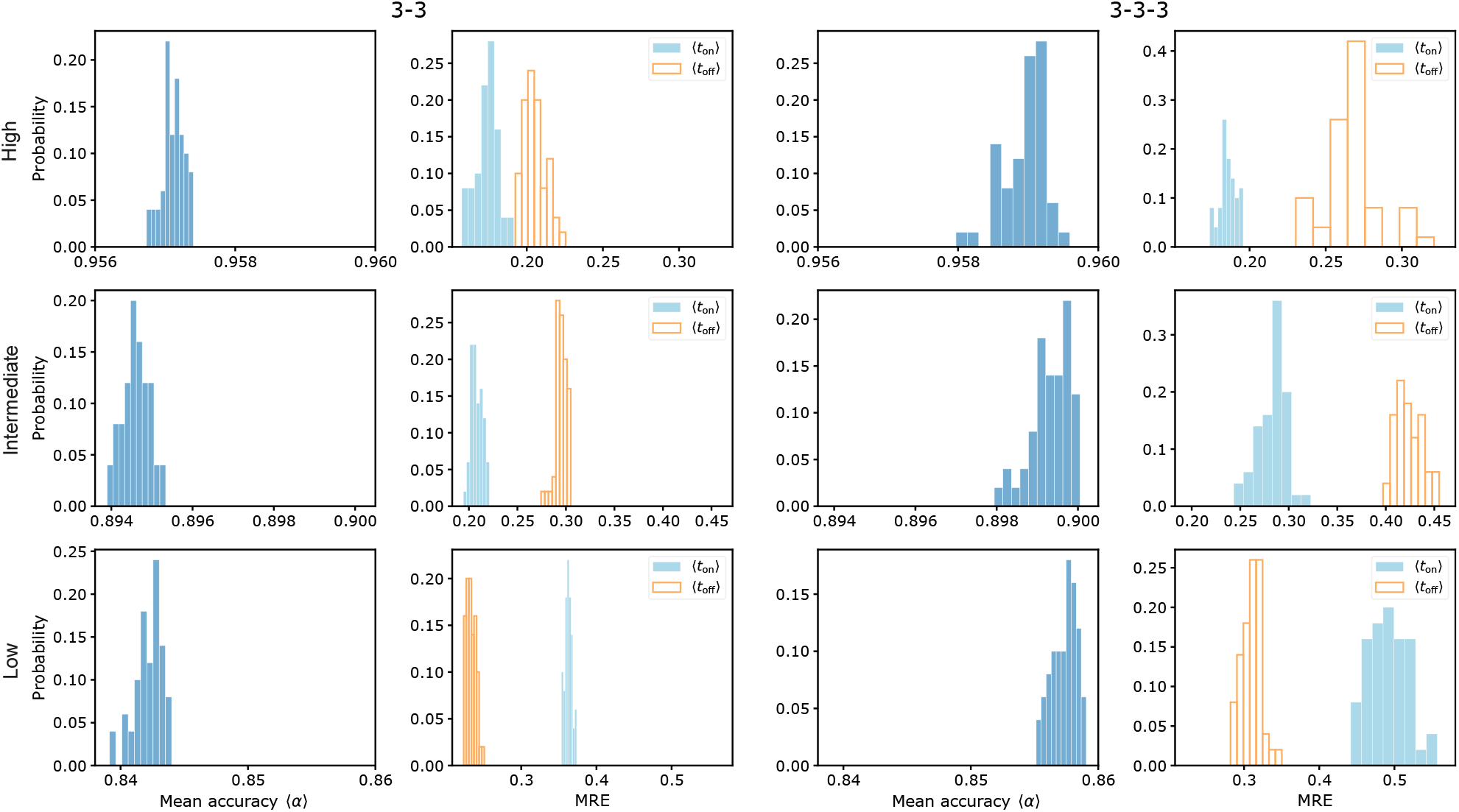
Comparison of the performance of two candidate neural network structures for DART. Left panel: Two convolutional layers, each with kernel size of 3 (denoted as 3-3). Right panel: Three convolutional layers, each with kernel size of 3 (denoted as 3-3-3). In both panels, the first column shows the posterior distribution of the mean binary vector accuracy ⟨*α*⟩ over 50 random seeds, and the second column shows the posterior distribution of the MRE for the estimated mean on- and off-times, also over 50 random seeds (see Supplementary Text S4 for details). Each row corresponds to a different burstiness level, decreasing from high to low from top to bottom (for definitions of the burstiness levels, see Materials and Methods, Section 4.2.2).

